# Coordination and persistence of aggressive visual communication in Siamese fighting fish

**DOI:** 10.1101/2024.04.29.591330

**Authors:** Claire P. Everett, Amy L. Norovich, Jessica E. Burke, Matthew R. Whiteway, Pei-Yin Shih, Yuyang Zhu, Liam Paninski, Andres Bendesky

## Abstract

Animals coordinate their behavior with each other during both cooperative and agonistic social interactions. Such coordination often adopts the form of “turn taking”, in which the interactive partners alternate the performance of a behavior. Apart from acoustic communication, how turn taking between animals is coordinated is not well understood. Furthermore, the neural substrates that regulate persistence in engaging in social interactions are poorly studied. Here, we use Siamese fighting fish (*Betta splendens*), to study visually-driven turn-taking aggressive behavior. Using encounters with conspecifics and with animations, we characterize the dynamic visual features of an opponent and the behavioral sequences that drive turn taking. Through a brain-wide screen of neuronal activity during coordinated and persistent aggressive behavior, followed by targeted brain lesions, we find that the caudal portion of the dorsomedial telencephalon, an amygdala-like region, promotes persistent participation in aggressive interactions, yet is not necessary for coordination. Our work highlights how dynamic visual cues shape the rhythm of social interactions at multiple timescales, and points to the pallial amygdala as a region controlling engagement in such interactions. These results suggest an evolutionarily conserved role of the vertebrate pallial amygdala in regulating the persistence of emotional states.

## Introduction

The coordination of behavior among individuals is a fundamental feature of social interactions^1,2^. Such coordination can positively correlate the behaviors of two or more individuals in space and time, as seen during fish schooling and bird flocking, and also during competitive encounters such as head butting in rams^3,4^. In other instances, animals anticorrelate their behavior, such that “turn taking” occurs. A canonical example of turn taking is acoustic communication, displayed by many invertebrate and vertebrate species in both cooperative and agonistic interactions^5–7^. Turn taking in agonistic interactions often involves alternating attack with defense, as seen in the ritualized exchange of strikes during territorial conflicts in mantis shrimp^8,9^, in chasing and fleeing in stickleback fish^10^, and in human martial arts.

How animals determine when it is their turn to perform a behavior is best understood in acoustic interactions. In singing mice, for example, the end of a singing bout triggers the start of singing by a partner^5^. The psychophysics, behavioral dynamics, and neuronal mechanisms of turn taking reliant on sensory modalities other than audition, however, are not well understood. Part of the difficulty in characterizing the precise sensory cues that shape turn-taking behavior stems from the fast dynamics and highly correlated nature of multiple behavioral features, both within and across sensory modalities, and within and across the interlocutors. Furthermore, the neural substrates involved in the sensorimotor transformations that support turn taking and in sustained participation in a social interaction, are not well known.

To study visually-driven turn-taking behavior, we focus on the Siamese fighting fish (*Betta splendens*, or simply ‘betta’). Betta is a particularly aggressive species of fresh-water fish endemic to Thailand, which has been used for centuries in organized fights^11,12^. Betta aggressive encounters start with a visual display phase, in which animals take turns at facing while flaring their opercular gill covers, alternating with turning sideways while spreading their fins and sometimes beating their tail^13^. This display phase can end when one individual —usually the one who flares less— surrenders and flees^13–16^, or can escalate into an injurious phase involving biting^13,14,17–19^. In this study, we dissect the dynamic interaction sequences and relevant visual cues through quantitative analyses of aggressive behavior of fish interacting with a conspecific or computer animations. We discover the visual features that shape turn-taking behavior acutely and that promote continuous participation in the display phase. We then identify the brain regions that are particularly active while fish display aggression and that promote continuous engagement in the aggressive display.

## Results

### Betta take turns during aggressive displays

Turn-taking aggressive displays have been noted in betta located in adjacent transparent tanks^13^. However, whether turn taking occurs in more ethological settings —when individuals have physical access to each other— has not been documented. We placed a pair of male betta fish in the same tank and measured flaring of the opercular gill covers (henceforth ‘flaring’), tail beating, and biting behaviors (**Fig 1A,B**). The first flaring event occurred 23 seconds after fish obtained physical access to each other, whereas the first tail beating occurred 34 seconds after the first flaring. Each flaring bout lasted an average of 1.1 seconds (±0.99 seconds standard deviation) and rarely overlapped with flaring by the opponent (p=10^-48^ by Fisher’s exact test for lack of overlap; r=-0.94). During the first four minutes of the interaction, flaring occurred 18% of the time and tail beating occurred much less frequently —only 2% of the time. The first bite was seen at 3.6 minutes into the interaction, and a clear transition into biting by both fish occurred only 30 seconds later. At that point, biting was the predominant behavior, while flaring and tail beating essentially disappeared. Altogether, these patterns are consistent with aggressive behavior in betta going through a display phase, when animals take turns at flaring, followed by a phase of contact fighting where biting replaces flaring.

**Figure 1:**
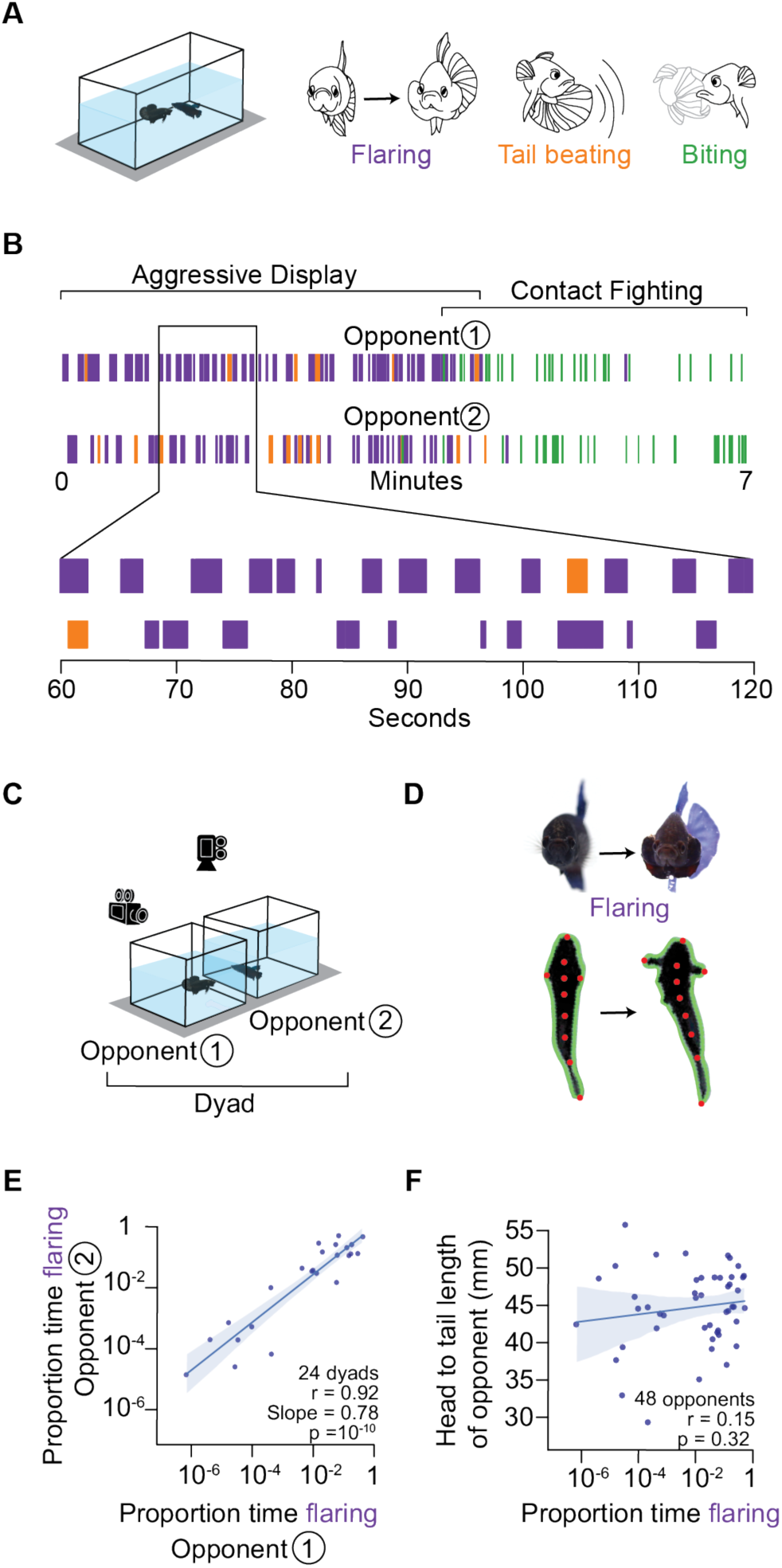
Betta take turns during aggressive displays and scale their aggressive response to match that of their opponent. **(A)** Opponents were placed in the same tank and multiple aggressive behaviors (flaring, tail beating, and biting) were scored and shown as a raster plot in **(B)**. **(C)** Behavioral paradigm consisting of a dyad in neighboring tanks, video recorded from the top and side. **(D)** Fish keypoints (red dots) and contour (green outline) are tracked to quantify behavior. **(E)** Correlation of the proportion of time flaring between randomly chosen opponents 1 and 2. Each point denotes a dyad. **(F)** Correlation of proportion time flaring of either opponent 1 or 2 with the average head to tail length of their respective opponent.

The contactless turn-taking aggressive displays of betta can be elicited by visual cues alone^14,20–25^ and this is often achieved using a mirror ^23,24,26^. However, this mirror setup is not conducive to studying turn taking, since fish only see a simultaneous reflection of their own behavior. Instead, we examined turn taking behavior by placing two fish in adjacent tanks while we videorecorded them from the top and side (**Fig 1C**). To automatically identify flaring behavior, we tracked the contour and multiple keypoints of each fish^27^ (**Fig 1D**). We then trained a supervised behavioral classification model^28^ to automatically score flaring behavior. The model leveraged the tracked anatomical elements, along with geometrical features derived from these elements, such as the angle formed between the tip of the nose and the tips of the operculum and the orientation of the fish, as well as basic movement behaviors including speed and turning angle (**SFig 1A-G**). This analytical procedure achieved an F1 score (reflecting both precision and recall using manual scoring as the ground truth) of 0.75 (**SFig 1I**). When we required even higher sensitivity and specificity, we complemented the automated method with manual scoring, and we note these cases in the Methods.

Using these behavioral and analytical methods, we found that the fraction of time spent flaring by two randomly-paired opponents across tanks was highly correlated (r=0.92, p=10^-10^; **Fig 1E**), consistent with previous findings^13^. In contrast, the fraction of time spent flaring was not correlated with the size of an opponent (r=0.15, p=0.32; **Fig 1F**). These observations indicate that dynamic visual cues —rather than size— are sufficient for betta to scale their aggression to match that of their opponent.

### Two complementary paradigms to study visually-evoked aggressive displays

To discover the visual cues that shape turn-taking aggressive displays, we studied betta behavior in two complementary paradigms: (1) exposing individuals to another betta (a “conspecific”) in an adjacent transparent tank and (2) exposing individuals to a naturalistic computer animation of a betta performing an aggressive display (**Fig 2A,B**; see Methods). The first paradigm captures a palette of natural behaviors, whereas the second has the advantage of presenting consistent and repeatable stimuli to all fish. The animation was created by tracking a betta performing an aggressive display to a conspecific —with intervals of flaring, as well as changes in orientation, velocity, and position— and then latching this real movement pattern onto the 3D-lattice of a realistic-looking betta (**SVideos 1,2,3** and Methods). The animation is 37 seconds long and was shown for a total of 10 minutes (looped 16.2 times) to each individual. Fish are exposed to both paradigms in a random order, separated by a 10-minute break. Prior to revealing the stimulus, we found that animals spend only ∼30% of the time in the quarter of the tank closest to the stimulus side (the other conspecific’s tank or the monitor), but ∼60% of the time after reveal (p=10^-9^; **Fig 2C,D**). Furthermore, in both paradigms animals spend only ∼50% of the time facing the stimulus side before reveal, but ∼68–80% after reveal (p=10^-14^; **Fig 2E,F**). We found no significant difference in time spent near the stimulus or time facing the stimulus between real conspecific stimulus and the animation (stimulus × condition p=0.67 and p=0.18, respectively). In both paradigms, fish flared exclusively after the stimulus was revealed (exposure p=10^-23^), consistent with flaring requiring visual cues. Upon stimulus reveal, fish flared less towards a conspecific than to the animation but flaring against the animation decayed faster over time (**Fig 2G**). These patterns led to a similar proportion of time spent flaring in both paradigms during the 10-minute stimulus exposure period: ∼22% against a conspecific and ∼30% against the animation (stimulus × condition p=0.08; **Fig 2H**). Altogether, the results indicate that fish engage with both types of stimuli to a similar extent and that the naturalistic animation is as effective as real conspecifics at eliciting aggressive displays from an opponent.

**Figure 2:**
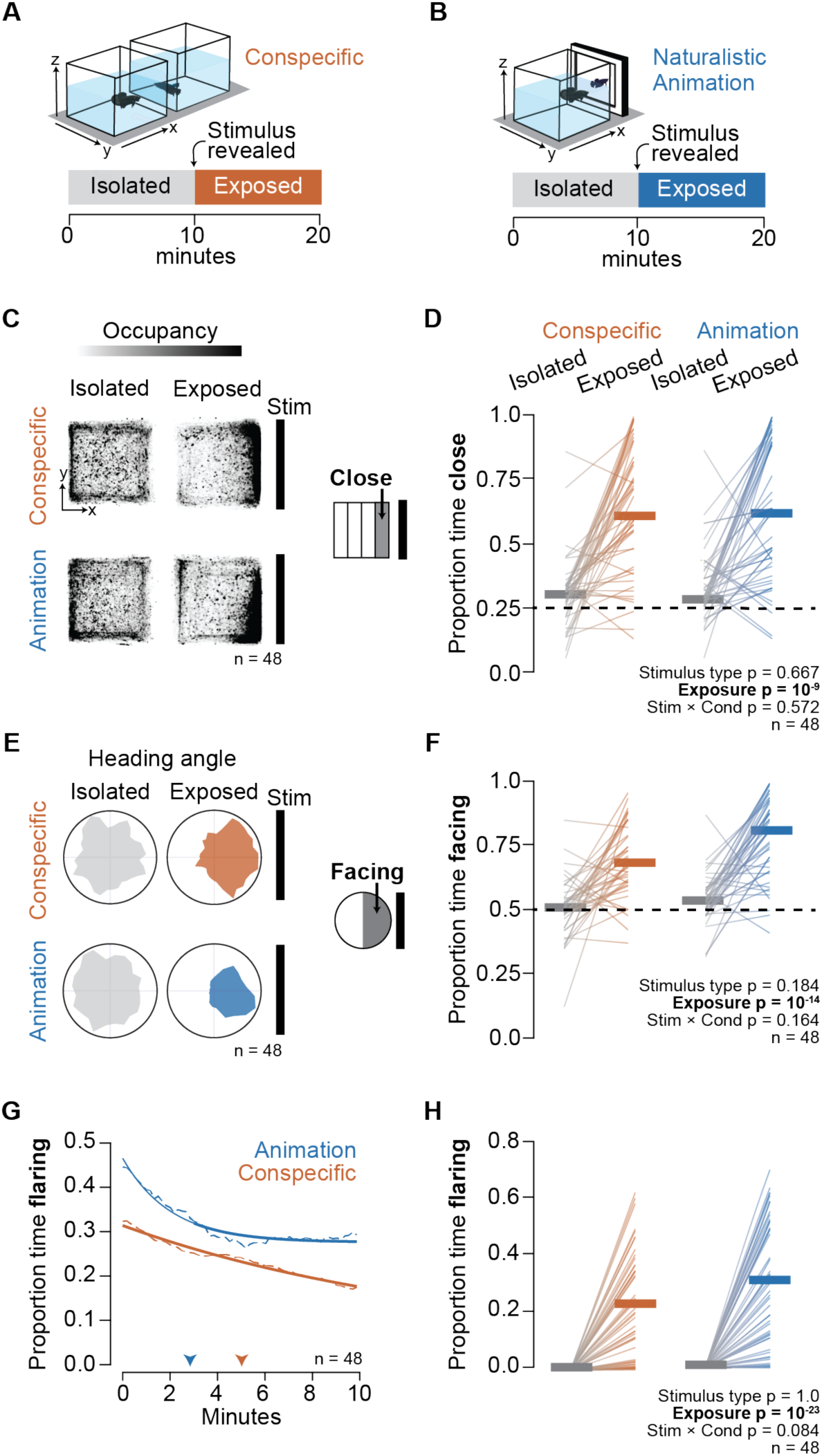
Naturalistic animations evoke as strong a flaring response as a conspecific. **(A)** Conspecific paradigm: two opponents in neighboring tanks. **(B)** Naturalistic animation paradigm: one opponent faces a naturalistic animation modeled after a male displaying aggression (see Methods). **(C)** Heatmaps showing tank occupancy (from a top view) by fish during isolation and exposure. **(D)** Proportion time spent in the quarter of the tank closest to the stimulus. Dotted line denotes chance level. **(E)** Polar plots showing distributions of the head orientation during isolation and exposure. **(F)** Proportion time spent facing within 180° of the stimulus. Dotted line denotes chance level. **(G)** Average persistence of proportion time flaring over the course of the exposure period (dashed-lines) with exponential curve fit (solid-lines). Arrow heads point to the time when the proportion time flaring decreases to 0.75× of max. **(H)** Proportion time flaring during isolation and exposure. **(D,F,H)** Thin lines denote individuals, horizontal thick lines denote the median. p-values by mixed models ANOVA.

### The visual cues that modulate betta aggressive behavior

We took advantage of the repetitive nature of the animation —presentation in a loop 16.2 times— to determine how different parts of the animation are related with flaring responses of the observer. Notably, fish synchronize their flaring behavior to defined epochs of the animation (**Fig 3A,B**). Qualitatively, fish flare the least when the animated fish is facing forward, and most when the animated fish is lateral, and they demonstrate this pattern repeatedly across loops of the animation. Overall, 81% of fish (17/21) synchronized their flaring (see Methods, **SFig 2**) across loops of the animation (**Fig 3C**). To quantitatively relate flaring behavior to specific features of the animation, we measured the correlation between flaring and multiple dynamic features of the animation. These features included position, speed, velocities in different directions, acceleration, and elevation, as well as orientation and flaring (**SFig 3**). The strongest correlations were with flaring (r=-0.55), lateral orientation (r=0.3), and elevation (r=0.2) (**Fig 3D**). The anticorrelation with flaring is consistent with the turn-taking behavior observed when two fish have physical access to each other (**Fig 1B**). Flaring occurs almost exclusively when betta face their opponent and not when they are lateral (**SFig 4**). Therefore, the correlation between flaring and a lateral orientation of the animated fish is also consistent with turn taking. The correlation between flaring and the elevation of the animated fish was surprising and we explore this in a later section. The correlations between the behavior of the animation and the flaring of the observer were remarkably consistent when measured against a real conspecific, indicating that the animation captured visual elements that naturally trigger a flaring response (**Fig 3E and SFig 3**).

**Figure 3:**
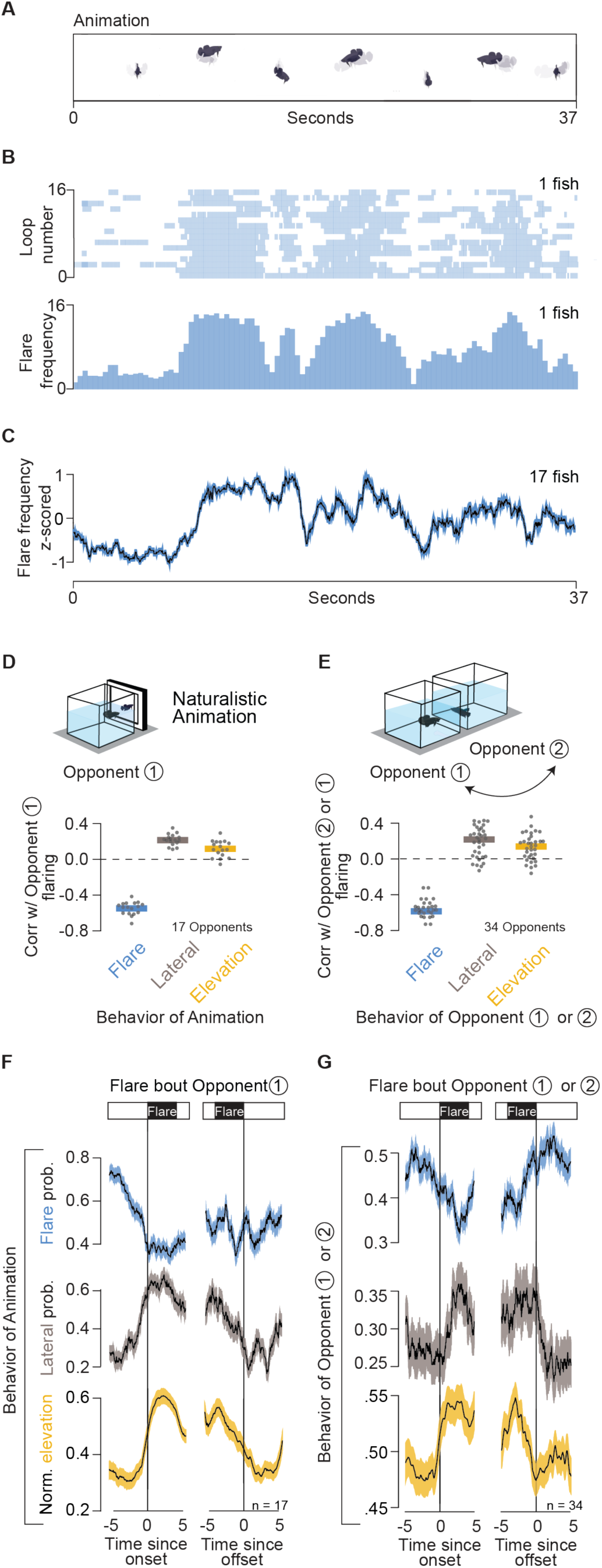
Betta coordinate their flare response with dynamic visual cues. **(A)** Selected frames from one loop of a naturalistic betta animation. **(B)** Raster plot (top) and histogram (bottom) showing flaring during each timepoint in the animation. **(C)** Z-scored flaring frequency (mean±SEM). **(D)** Correlation between flaring of opponent 1 with behaviors of the animation. **(E)** Correlation between flaring of opponent 1 or 2 with behaviors of their opponent. (**F, G)** Peri-event time histograms (mean±SEM) of changes in flare probability, lateral orientation probability, and normalized elevation (see Methods) of the virtual fish **(F)** or real opponent **(G)** aligned to either the onset (left) or offset (right) of flaring bout of the opponent.

To parse the correlations between stimulus and flaring response in a more temporally precise manner, we generated peri-event time histograms aligned to the beginning or end of a flaring event (**Fig 3F,G**). This allowed us to characterize which behaviors, either of the animation or of the real opponent, may cause flaring to start or to stop. We found that fish began to flare when the animated fish stopped flaring, went lateral, and elevated itself. By contrast, fish stopped flaring as the animation turned from lateral to facing forward, and as it descended on the screen. Interestingly, fish did not time the end of a flare with the start or end of a flare from the animation, suggesting that observing a change in orientation from lateral to facing and a decrease in elevation are sufficient to interrupt flaring. Interactions between real-opponents led to qualitatively similar observations (**Fig 3G**), though interpretations are more difficult because interacting fish respond dynamically to each other, whereas fish cannot influence the behavior of the animation. Overall, three fourths of flaring bouts start when the opponent has started to close its operculum or has stopped flaring altogether. Results from quantitative analysis of aggressive encounters with conspecifics and animations suggest that both the onset and offset of a flaring event are regulated by specific visual cues; that animals start flaring when their opponent turns lateral, stops flaring and elevates itself; and that they stop flaring spontaneously or when their opponent turns to face them and moves downward in the water column.

### Role of flaring in the aggressive display

When betta start to flare, they also curve their body as they turn to face their opponent, transiently increase their speed, and lower themselves in the water column (**SFig 4**). It is therefore challenging to know which of these visual features shape the flaring response of the opponent. To decouple flaring from facing, we created two animations in which a fish alternates between two phases: (1) facing forward while stationary and (2) turning lateral and swimming to the other side of the screen, in a loop (**Fig 4A,B** and **SVideos 4,5**). The two animations differ only in whether fish flare while facing forward. Individuals were exposed to both animations (each in a 10-minute block) in a balanced order. The onset of flaring bouts reached its maximum when the animated fish turned lateral, and its minimum when the animation faced forward (**Fig 4C**). Conversely, offsets of flare bouts reached a maximum when the animated fish faced forward (**SFig 5**). Surprisingly, this pattern was indistinguishable regardless of whether the animated fish flared or not. Consistent with the time histograms (**Fig 4C**), a linear model for flaring onset was significantly affected by the phase of the animation (p=10^-6^), but not by the type of animation (flaring or not flaring; p=0.47), nor by an interaction between animation phase and type (p=0.29; **Fig 4E**). Notably, although flaring by the stimulus did not acutely affect flaring by the observer, it did lead to a 41% increased (p=0.014) median time flaring during the 10 minutes of the animation (**Fig 4D**). This increase in total flaring time was caused in part by a more persistent flaring across the 10 minutes of exposure (**Fig 4F**). These results indicate that betta can modulate their flaring response from watching an opponent change in orientation between lateral and forward facing and that flaring is not required for turn taking behavior. Instead, flaring seems to encourage continued engagement in aggressive interactions.

**Figure 4:**
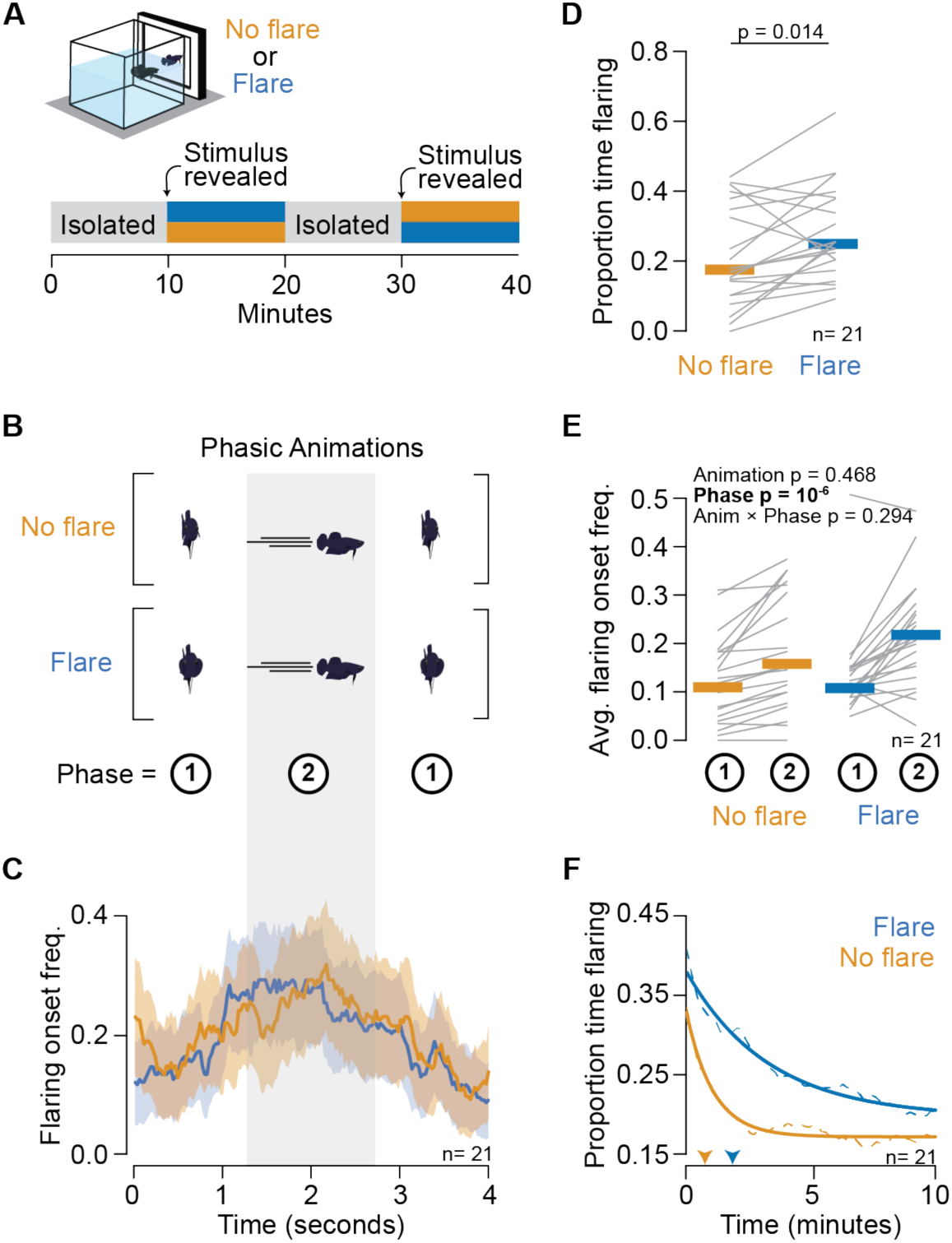
Flaring of a stimulus is not necessary for the synchronization of flaring but promotes a more persistent flare response. **(A)** Behavior set up and paradigm presenting two phasic animation types (No Flare or Flare) to fish in a balanced order. **(B)** Animations were composed of two alternating phases with distinct combination of flare, orientation, and speed state. **(C)** Frequency of flaring bout onsets aligned to a loop of the animation. **(D)** Proportion time flaring against each animation. **(E)** Average flaring onset frequency for each phase (1 or 2) in each animation. **(D,E)** Thin gray lines denote individuals, horizontal thick lines denote the median. p-values by **(D)** two-tailed paired t-test or **(E)** mixed methods ANOVA. **(F)** Average persistence of proportion time flaring over the course of the exposure period (dashed-lines) with exponential curve fit (solid-lines). Arrow heads point to the time when the proportion time flaring decreases to 0.75× of max.

### Role of elevation in the aggressive display

Our finding that a higher elevation of the opponent promotes a flaring response (**Fig 3D-G**) was unexpected, as this has not been described as a relevant visual cue for aggressive behavior in betta nor in other aquatic animals. Elevation in the water column, however, can have important ethological relevance since many elements of the life and behavior of betta biology occur near the surface. As anabantoids with a lung-like labyrinth organ, betta can breathe air^29^. During fights, betta increase oxygen consumption and the rate of ascension to the water surface to breathe^24,26,30^. Furthermore, betta males build bubble nests at the water surface and show aggressive territorial behavior near their nests^13,29,30^. Therefore, we hypothesized that animals display more aggression (a sign of territoriality) the closer their opponent is to the surface, rather than the farther it is from the bottom. To test this hypothesis, we exposed fish to an animation presented at varying distances from the top and from the bottom, decoupling the two by also varying the height of the water (**Fig 5A,B**). The results strongly supported the hypothesis: fish flared the most to stimuli that appeared closest to the top of the water column and least to stimuli closest to the tank floor (p=10^−4^ for distance from top and p=0.25 for distance from bottom in a generalized linear model). Notably, when presented at the top of the water column, the simple phasic animation that alternates between facing and flaring, with turning laterally and moving to the other side (which we used as the “Flare” condition in **Fig 4**), evokes as much flaring from the observer as the more complex naturalistic animation (**Fig 5C**). This attests to the strength of a simple stimulus in eliciting aggression, if presented close to the surface.

**Figure 5:**
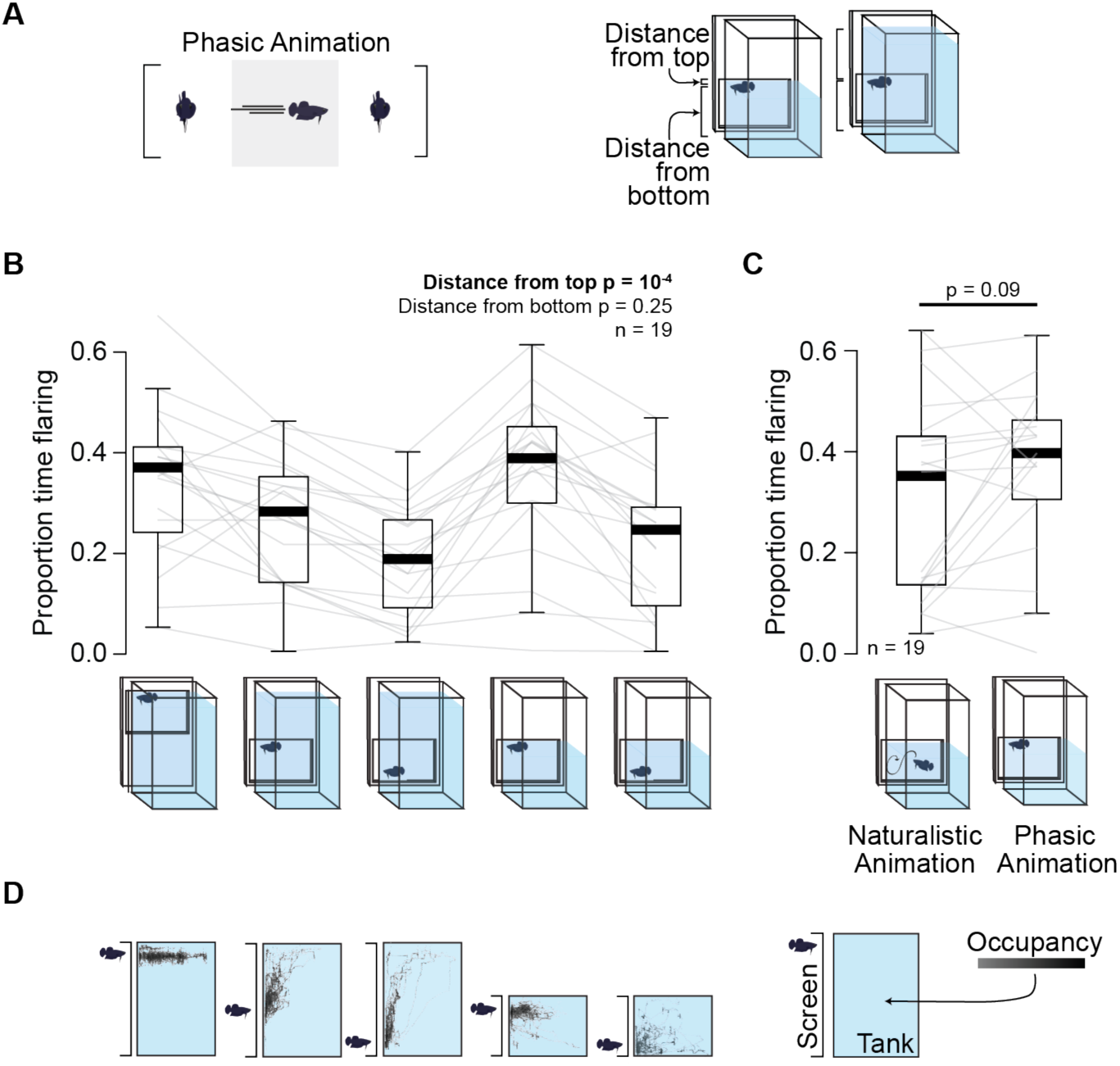
Flaring increases when opponent is close to the surface. **(A)** Left: Side view of a tank showing position of screen. Middle: Phasic animation in which stimulus maintains its elevation. Right: animation was shown at varying distances from surface and bottom of water by varying the elevation of the animation and the height of the water. **(B)** Proportion time flaring against stimuli with varying distances from surface and bottom. **(C)** Comparison of proportion time flaring against naturalistic animation versus phasic animation at elevation closest to the surface. **(D)** Heatmap of exemplar individual betta against each elevation condition.

We further explored how betta adjust their own elevation to interact with stimuli at different depths. When presented with the animations that do not change in depth, animals aligned themselves with the stimuli (**Fig 5D**). However, this alignment was more variable the farther the stimuli were from the surface. This might reflect a stronger drive to interact aggressively with opponents near the surface, difficulty in maintaining the buoyancy necessary to go to the bottom (since betta take in air more frequently when they fight), or spending less time close to the bottom due to the frequent trips to the surface to breathe. In encounters between two real animals, before gaining visual access to each other, there was a bimodal distribution of elevation, with animals spending most of their time close to the bottom or close to the surface and least time in between (**SFig 6**). After exposure to each other, however, both animals spent most of their time close to the top (**SFig 6**) suggesting a preference for elevated position during displays.

### The caudal dorsomedial telencephalon promotes engagement in aggression

Next, we aimed to determine which brain regions modulate the aggressive display. To that end, we began by performing a brain-wide screen for neurons that were active when fish were engaged in an aggressive display (**Fig 6A,B** and **SFig 7**). We used phosphorylation of the ribosomal S6 protein (pS6) as a proxy for neuronal activity during display^31,32^. Four out of the 28 brain regions assessed had a significant (p<0.05) increase in the number of pS6+ neurons in displaying animals compared to unexposed controls (**SFig 7**). No region showed a decrease in activity during display. The four regions with increased activity were the caudal dorsal medial area of the pallium (cDm; 2.3× increase in pS6+ cells, p=0.003), the supracommissural (Vs; 1.5×, p=0.026) and postcommisural (Vp; 2.3×, p=0.029) nuclei of the ventral telencephalon, and the caudal preglomerular nucleus (PGc; 3.1×, p=0.017). Of these, cDm has previously been suggested to promote aggression in betta^18^ and promote arousal and defensive behaviors in several other teleost fishes^33–35^. Based on its development and cytoarchitecture, Dm is considered homologous to the tetrapod amygdala^36–39^. Thus, cDm stood out as a promising candidate for involvement in the modulation of visually-evoked aggression.

**Figure 6:**
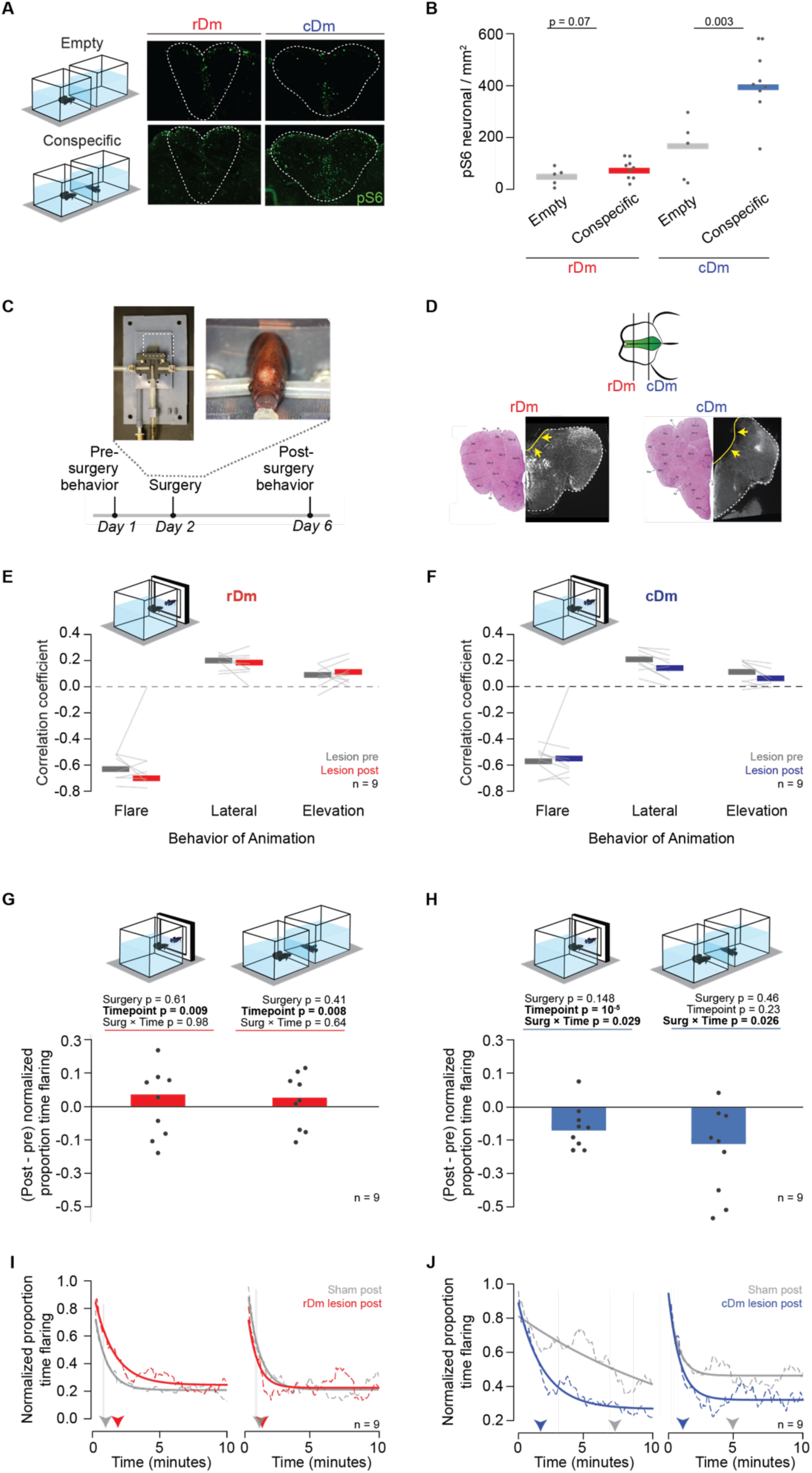
Caudal Dm promotes persistent engagement in an aggressive interaction. **(A)** Left: Betta were exposed to either a conspecific or empty tank for a brain-wide activity screen. Right: Example sections of rDm and cDm with pS6+ cells. **(B)** Quantification of pS6+ cells in rDm and cDm following conspecific or empty tank exposure. p-values by Welch’s t-test. For other brain regions see Suppl. Fig. 7. **(C)** Custom survival surgery set up for Dm lesioning and experimental timeline. **(D)** Example sections showing targeted lesions of rDm and cDm. Arrows point to the brain region removed by the lesion. **(E, F)** Correlation between flaring of **(E)** rDm or **(F)** cDm lesioned fish pre- and post-surgery with behaviors of the naturalistic animation. **(G, H)** Change in the proportion time flaring towards animation (left) and conspecific (right) following lesion of **(G)** rDm or **(H)** cDm, normalized by change in proportion time flaring of sham-operated individuals. p-values by mixed-methods ANOVA. **(I, J)** Normalized average persistence of time flaring over the course of the exposure (left: animation; right: conspecific) period (dashed-lines) with exponential curve fit (solid-lines), normalized to the maximum flaring of each individual. Arrow heads point to the time when the proportion time flaring decreases to 0.5× of max following **(I)** sham or rDm lesions, and **(J)** sham or cDm lesions.

Teleost Dm has been implicated in sensory processing, including in vision^35,40^. To examine whether Dm received direct visual input in betta, we performed anterograde tracing from the retina using cholera toxin b (ctb), which is transported anterogradely in retinal ganglion cells (RGCs)^41–43^. Consistent with findings in other teleost fish^44,45^, we observed ctb+ RGC terminals in the optic tectum and in a cluster of pretectal nuclei, but not in Dm (**SFig 8**). This indicates that Dm does not receive direct input from the retina. Instead cDm might receive visual information via ascending visual pathways to the telencephalon, as has been seen in other teleost fish^46^.

We next directly examined the role of cDm in aggression by lesioning cDm and subjecting these animals to our controlled behavioral assays (**Fig 6C-J**). A previous study lesioned cDm in betta, but it did not report the extent, nor accuracy of the lesions, nor any controls^18^. We bilaterally ablated this region by aspiration using a novel setup for betta stereotaxic surgeries (**Fig 6C,D**). To control for a potential effect of the surgery on aggressive display behavior, we sham-lesioned animals by putting them through the same surgical procedure, including anesthesia and craniotomy, but without lesioning their brain. As an additional comparison with cDm lesions, we also lesioned the rostral Dm (rDm), which did not show a significant increase in pS6 labeling following during aggressive displays (1.1× relative to unexposed fish, p=0.07; **Fig 6B**). The behavior of experimental animals was quantified before and after surgery to enable more powerful within-subjects comparisons. We let animals recover for four days after surgery, minimizing the opportunity for plasticity post-lesion, which can complicate the interpretation of the effect of lesions^47^. After behavioral testing, four lesions (16.7% of lesioned fish) were discarded.

Compared to controls, lesions did not significantly alter swimming speed, the proportion of time spent facing or in close proximity to the stimulus, or the number or duration of flaring bouts (**SFig 10** and **SFig 11**). We next evaluated whether rDm or cDm might be involved in the synchronization of displays with the naturalistic animation. Neither rDm nor cDm lesions affected the timing of flaring relative to the animation (**Fig 6 E,F** and **SFig 12**). However, cDm lesions, but not rDm lesions, reduced the proportion of time flaring towards both the naturalistic animation and a real opponent (**Fig 6 G,H**). This reduced flaring of cDm-lesioned animals was mostly a consequence of decreasing participation in the aggressive display over time: Animals with cDm lesions began flaring at similar levels as sham-lesioned animals, but their flaring rate decayed faster over the next 10 minutes (**Fig 6 I,J**). In contrast, rDm lesions did not affect the trajectory of flaring rates, compared to sham-lesioned controls. Thus, cDm is necessary for promoting continued engagement in aggressive interactions.

## Discussion

By studying visual encounters with conspecifics and with controlled animations, we discover the key features that shape turn-taking aggressive behavior in betta. When fish 1 stops flaring and facing and begins to reorient laterally, that triggers facing and flaring by fish 2; fish 2 then stops flaring and turns lateral, inducing facing and flaring by fish 1, leading to turn-taking cycles. Sometimes fish don’t wait for “their turn” and instead start facing their opponent and flaring while the opponent is still flaring, which induces the opponent to stop flaring and turn sideways. Therefore, betta usually, but not always, wait for the other fish to stop flaring before flaring themselves, similar to the dynamical turn taking observed in vocal communication in humans and other animals. This implies that a similar neuronal circuit logic may gate sensorimotor transformations in both acoustic and visual communication^48^.

Because fish flare almost exclusively when facing their opponent, flaring and orientation are highly correlated. By presenting fish with animations that decouple these behaviors, we find that forward facing without flaring is enough to suppress flaring from the opponent. Notably, although flaring is not required to acutely modulate flaring by the opponent, it is required to promote continued engagement in aggressive display by the opponent over a timescale of minutes. Therefore, some visual cues shape social interactions acutely and others do so chronically.

We find that the caudal portion of the dorsomedial telencephalon (cDm) is differentially active while animals display aggression. cDm does not appear to be involved in the sensorimotor transformations required for turn taking. Instead, cDm promotes continuous engagement in an aggressive display with an opponent. Dm shares developmental, cytoarchitectonic, and connectivity features with the tetrapod pallial amygdala^35^. Our findings indicate that in addition to developmental and anatomical homology, fish cDm, like the mammalian amygdala, is also involved in maintaining persistent behavioral states related to emotion and self-preservation ^49,50^. This indicates that the amygdala has an evolutionarily conserved role across vertebrates in regulating behavioral states.

Altogether, we discover how select social visual cues set the rhythm of social turn-taking behaviors while others modulate the duration of social interactions. Further, we find that an amygdala-like region is an important node for maintaining a persistent involvement in social interactions.

## Acknowledgments

Michael Long advised on PDLC screens for behavioral testing. Miles Marshall developed an early version of an animated betta. Taiga Abe helped implement pose estimation. Madison Lichak and Alec Palmiotti provided technical support. Sarah Aktari, Christina Lin, Jane Chen, Bliss Bagnato-Conlin, Valentine Andreu and Clara Liff helped with experiments. Tanya Tabachnik aided in the 3D printing of stereotaxic platform. Luke Hammond and Darcy Peterka provided training and advice on microscope imaging. This work was supported by the following grants to A.B.: Searle Scholarship, Sloan Fellowship in Neuroscience, and National Institutes of Health (NIH) R34NS116734, and R35GM14305; to C.E.: Vision Sciences Training Grant T32EY013933. A.L.N.: the Simons Society of Fellows Junior Fellowship 855220; to J.E.B.: National Science Foundation (NSF) Graduate Research Fellowship DGE-2036197, P.-Y.S.: NIH K99GM151689, L.P.: NSF 1707398, Gatsby Charitable Foundation GAT3708, Simons Foundation 543023, NIH U19NS123716.

## Author contributions

Conceptualization, C.P.E and A.B.; Methodology, C.P.E., A.L.N., J.E.B., Y.Z., M.W.; Validation, C.P.E; Formal Analysis, C.P.E, A.L.N., Y.Z., M.W.; Investigation, C.P.E., A.L.N., J.E.B, P.-Y.S; Writing – Original Draft, C.P.E and A.B.; Writing – Review & Editing, C.P.E, A.B., A.L.N, J.E.B, M.W., L.P.; Visualization, C.P.E. and A.B., Supervision, A.B.; Funding Acquisition, A.B.

## Materials and methods

### Experimental Model and Subject Details

Adult male betta bred for fighting were used for all experiments including behavior. Fighting males were sourced from an independent breeder in Bangkok, Thailand or then bred in-house. Ornamental males were only used for pS6 activity mapping and were either obtained from Petco or bred in-house. All males were 6–18 months of age when tested. Betta were maintained under standard husbandry and housing conditions^51^. Adult betta were individually housed and visually-isolated from other fish for at minimum two months before they were tested. All animal work was approved by the Columbia University Animal Care and Use Committee.

### Behavior Methods

#### Behavior Arenas

Closed behavioral arenas were constructed to ensure interactions with conspecifics and animations took place in a controlled environment devoid of visual disruptions. Arenas were constructed using 80/20 metal framing and white featureless foamboard walls. Each arena was 54 cm x 43 cm x 63 cm high. Waterproof string white LED lighting (SUPERNIGHT / 7000k color temperature) lining the top of the walls was used to illuminate the arena. The room that housed behavior arenas was kept at 26 °C. Each behavior arena had a top and side Raspberry Pi 3 Model B+ cameras with Raspberry Pi NoIR Camera Module V2 lenses. Cameras ran at 40 frames per second at a resolution of 1280×720 pixels. Tanks sat on platforms latched into laser cut stands to ensure a uniform distance of 1 inch between tanks or between tank and monitor.

#### Conspecific encounters

Individuals were placed inside a cubic tank (5×5×5 inches) and water was brought to a height of ∼4.5 inches. Tanks were separated by two Polymer Dispersed Liquid Crystal (PDLC) plates (12×6 inches, SW Store). These screens remain opaque during the habituation phase. At testing phase onset, current from Raspberry Pis is run through the PDLC screens causing them to become immediately transparent.

Fish were habituated in visual isolation from their opponent for 20 minutes. The last 10 minutes of habituation period were used to assess behavior during isolation. The PDLC screen then became transparent, and fish were given visual access to each other for the duration of a 10-minute test period.

### Animation encounters

Individuals were placed in a testing tank across from a monitor (4.6×7 inches LCD; ToGuard) controlled by a Raspberry Pi, displaying a white screen. At testing phase onset, the animations began playing in the monitor. Betta were habituated for 20 minutes followed by exposure to the animated stimuli. Exposure to an animation lasted for 10 minutes. If multiple animations were displayed to a fish, they were separated by 10-minute breaks in which the monitor displayed a white screen.

When testing the role of distance to the surface versus bottom, a single fish faced 8 animations (phasic animation at 5 different heights and naturalistic animation at 3 different heights). To avoid fatigue, animations were grouped into 3 different "clusters”. Each cluster contained 2–3 animations. Clusters were separated by 1-h breaks. Both the order of the clusters and animations within the clusters were randomized so that our results were not biased by potential priming or fatigue.

#### Designing animations

Animations were made using an open-source animation software, Blender v2.91. Naturalistic animations were crafted in two parts: 1) designing the 3D fish and 2) importing a naturalistic trajectory. The blender file is provided as SFile 1.

#### 3D fish

The volume of the virtual fish was built based on the body shape and size of a representative male. The fin shape and size were modeled after a photograph of a representative fighting male. The color was uniform across all parts of the fish (except the eyes, which were black) and was the average color value of a representative fighting male.

#### Trajectories

Naturalistic trajectories were created by tracking a male betta during an aggressive display using the motion tracking function in Blender (SVideo 1). The 3D-fish model was then latched to the track and the orientation and gill flaring were manually added. Phasic trajectories were crafted using custom python scripts and imported into Blender. The speed of the artificial paths (∼2 cm/sec) matches the average speed of 10 fish during their test periods in which they displayed towards visual stimuli.

#### Animation gill flare

The size of the operculum was changed manually. To optimize the ethological-relevance of flaring state in animations we measured change in width of a facing fish when the operculum was closed vs. open in a population of fish (n = 24). We then took the average change in width (1.3 cm) and used that to inform the full extension of the operculum in the animated fish.

### Quantifying Aggressive Display

This study utilized a mix of manual and automated scoring of aggressive behaviors. Automated scoring was used to assess the proportion of time spent flaring in a large population (n=48 fish, 96 trials), which required a high-throughput method (**Fig 1, 2**). However, when studying a smaller population or when scrutinizing subtle changes in flaring scale, synchrony, or onset/offset timing, manual scoring was employed (**Fig 3, 4, 5, 6**).

#### Manual scoring flaring bouts

Ground-truth flaring behavior was manually scored from the top-down video data using BORIS (Behavioral Observation Research Interactive Software)^52^. Flaring was divided into two sub-behaviors: partial flaring and full flaring. Partial flaring is defined as when the opercula are no longer touching the side of the body but not fully extended. Full flaring is when opercula reach and hold their maximum angle.

#### Automated scoring

Two DeepLabCut (DLC)^27^ networks were trained to track body parts from the video data. The first network tracked 18 body parts from the top view (1547 training frames from 20 individuals); the second network tracked 15 points from the side view (802 training frames from 15 individuals; **SFig. 1B**). Predicted markers with confidence values less than a threshold of 0.8 or a “jump” from a previous frame larger than 15 pixels were dropped, and linear interpolation was used to infill the missing data. Separately, the contours of the fish were computed from the raw video data using a custom OpenCV script (**SFig. 1C**; SFile 2). To fit a contour to fish, first the background was subtracted from each frame to create a binary image. The largest contour was found in the binarized image. The point in the contour closest to the DLC head and tail markers were considered the head and tail of the contour. Lines were fit to the 100 points surrounding the marked head and tail to determine the tail’s angle and how far it deviates from the body line. Behaviorally-relevant features were then extracted from the DLC markers (**SFig 1D)** and the contour (**SFig 1E**), including: orientation, speed, turning angle, head (x, y, z), centroid (x, y, z), tail (x, y), tail angle, tail deviation, and features of the operculum (operculum angle, left and right operculum angle and distance from spine). All features were smoothed using a median filter with a window of 11 frames (0.275 seconds). These behavioral features, and their associated labels from manual scoring, were then used to train a model that classifies the features into one of three available behavior classes for each time point: (1) no flaring; (2) partial flaring; (3) full flaring. The model is a dilated Temporal Convolutional Network (dTCN)^53^, which has shown good performance on similar behavioral classification tasks^28,54,55^ (**SFig 1F**).

#### dTCN architecture and loss

The dTCN model is composed of multiple “Dilation Blocks.” Each Dilation block is composed of two 1D convolution layers (filter size of 9-time steps and 32 channels per layer), each of which is followed by a leaky ReLU activation function and weight dropout with probability p = 0.1. An additional residual path bypasses the convolution layers and is combined with their output before the application of a final leaky ReLU function. The input features are passed through two such Dilation Blocks, and the output of the final Dilation Block is passed through a final dense layer followed by a softmax function to produce class probabilities. The model is trained using a standard cross-entropy loss function between the predicted class probabilities and the ground truth scores^56^. Because of the class imbalance present in this dataset, less frequent classes have higher weights in the loss function: if the total number of labeled frames is N, and the number of labeled frames for class *i* is N*_i_*, the loss for data points from class is weighted by N/N*_i_*.

#### dTCN training

Model training utilized 21 videos of individuals which contained a total of 1,512,000 hand scored frames (no flaring: 1,175,298 frames; half flaring: 177,978 frames; full flaring: 158,724 frames). These training videos were a mix of encounters with animations and conspecifics. The data were split into temporal sequences of 1,000 time points, and each training batch contained 8 such sequences. 90% of sequences were used for training, 10% for validation. Training terminated once the loss on validation data began to increase for 20 consecutive epochs; the epoch with the lowest validation loss was used for evaluation. All models were trained with the Adam optimizer using an initial learning rate of 10^-4^.

#### dTCN evaluation

Model evaluation utilized 19 videos of held-out individuals which contained a total of 1,368,000 hand scored frames. The argmax of the predicted class probabilities at each time point was used to construct ethograms of behavioral state (**SFig 1G,H**). To evaluate the models, the F1 score —the geometric mean of precision and accuracy— was computed for all behaviors across the 19 held-out test individuals (**SFig 1I**). Held-out test individuals were comprised of solely virtual fish encounters and therefore had overall higher flaring rates, compared to training set. Because F1 score is correlated with proportion time flaring, this lead to a higher average F1 score for withheld trials compared to training trials (**SFig 1J**).

A coarser measure of model performance related to the analysis in Fig 1 compares the proportion of time spent flaring (combining half and full flaring) predicted by the model and by hand scoring, for both training trials and held-out test trials (**SFig 1K,L**). All figures and supplemental figures visualize “combined” flaring, in which half and full flaring count as flaring.

### Statistical Methods

#### Synchronization towards animations

To measure synchronization, the flaring frequency (*X_S_*) at time (*t*) in the animation was calculated by adding flaring responses to each loop of the animation (*X_i_*) at time (*t*) and dividing by number of loops of the animation (*S*) (see equation 1).

The variance of the distribution of frequencies is then calculated. If a fish displays high flaring frequencies towards the animation at certain times (*t*) and low flaring frequencies at other times (*t*), instead of a uniform flaring frequency throughout, they are considered synchronized and will have a larger variance of frequencies 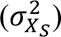 (see equation 2).

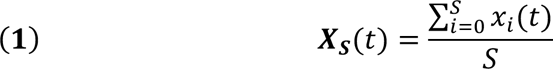

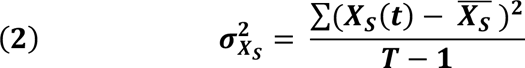

To determine whether fish are significantly synchronized to the animation, compared to chance, we performed a resampling of randomly offset controls. Flaring responses to each loop of the animation (*X_i_*) was shifted by a random value (θ) within the range of 0 to the length of the animation (*T*). With each flaring response randomly offset, we calculated the new variance of the frequencies 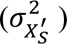 (see equation 3).

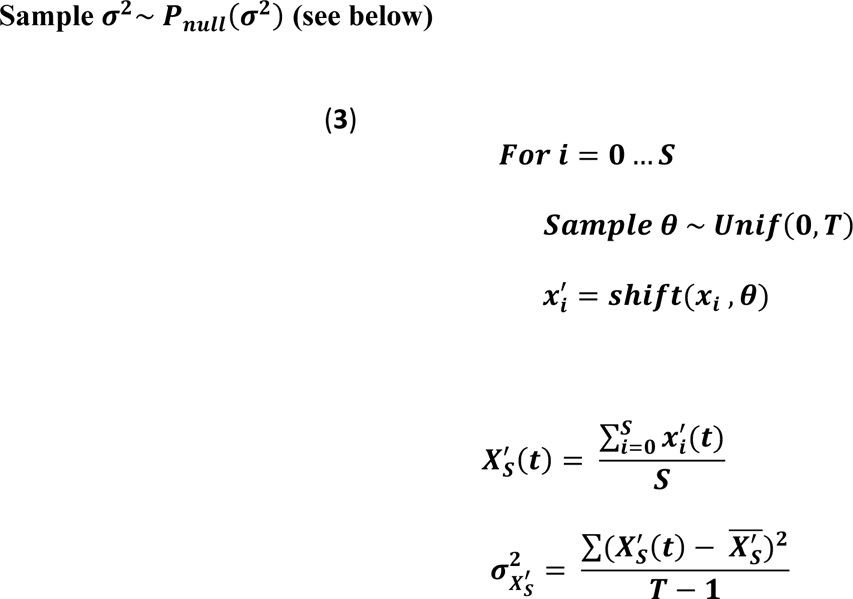

We repeated this resampling for 1,000 iterations to create a set of variances that can occur by chance while still maintaining the structure of the behavior. We calculated the 95^th^ percentile of the distribution of variances 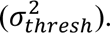 If the actual variance 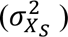 was greater than 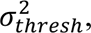 we concluded that the fish was significantly synchronized to the animation.

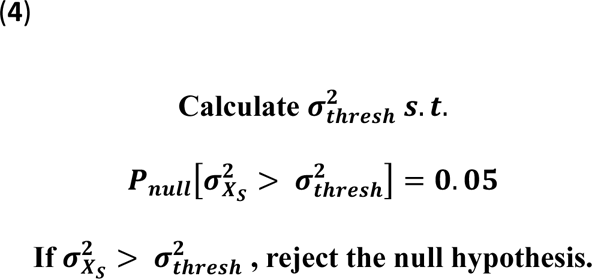

#### Correlations of flaring and stimulus behavior

Marker- and contour-derived features of shape and motion were gathered from both opponents during conspecific encounters, or for one opponent during animation encounters. Subsequently, we filtered time series data to include periods when either opponent was flaring during conspecific interactions, or when the opponent or animation was flaring during animation encounters. This step aimed to prevent artificially inflating high correlations of flaring observed in low-flaring dyads.

Next, we conducted point biserial correlations between flaring and opponent or animation features. In cases in which fish did not flare, we assigned a correlation value of 0 with all features. As a control, we rearranged the filtered time series of features in a random order and recalculated the correlation between flaring and each feature. A random shuffle opposed to a circular shift (as was done in quantifying synchronization towards animation above) was used since the correlation metric does not depend on temporal structure. We conducted paired *t*-tests between each population of actual versus shuffled correlation coefficients to assess the significance of the correlation.

#### Peri-event time histograms

We aligned the behavior of either the opponent fish or the animated stimulus with the onset or offset of a flaring bout exhibited by the focal opponent. Specifically, we chose flare bout onsets where a fish had not flared within the preceding 3 seconds and continued to flare for the subsequent 3 seconds. Similarly, for flare bout offsets, we selected instances where a fish had been flaring for 3 seconds prior and did not flare again for 3 seconds afterward. This selection criterion aimed to filter out flaring bouts that were either too brief or occurred in too close succession, minimizing potential noise in the opponent’s response.

To calculate the elevation PETH across time periods surrounding multiple bout onsets and offsets, we first conducted a normalization step on a period-by-period basis (**Fig 3F,G**). The elevation in the period surrounding each bout onset or offset was normalized by subtracting the minimum elevation within the period (3 seconds before and 3 after the onset or offset) and then dividing by the adjusted maximum elevation within that period. This normalization step accounted for absolute variations in elevation across different periods.

#### Flaring persistence

To quantify the proportion time flaring within a given sub-epoch within exposure period, the persistence of flaring for each trial was calculated by first binning the flaring time series into 8-second bins. A rolling mean window (width 0.357 seconds) was applied to the bins. The average persistence across individuals was calculated and an exponential curve was fit to the average proportion time flaring. The timepoint in which the fit exponential curve becomes ¾ (**Fig 2, 4**) or ½ (**Fig 6**) its maximum height is calculated.

### Histology, Circuit Tracing, and Ablation Methods

#### Activity- mapping

Betta were exposed to either a conspecific or empty tank for 30 minutes. Fish were euthanized 90 minutes following the end of exposure. Fish were euthanized by submerging ice water until immobile, followed by rapid decapitation. Heads were collected and fixed overnight in 4% paraformaldehyde (PFA) in 0.1M Phosphate buffer (PB) at 4 °C. Following fixation, brains were dissected out and washed 3× in phosphate buffer saline (PBS) before dehydration in 30% sucrose in 0.1M PB. Brains were embedded in OCT (Tissue Tek) for cryosectioning. 20 µm tissue sections were incubated overnight at 4 °C with primary antibodies diluted in 0.2% TritonX-100 PBS solution (0.2% PBT). Following 3× PBS washes, sections were incubated with secondary antibodies diluted in 0.2% PBT for 1 h at room temperature. NeuroTrace labeling (ThermoFisher; 1:200 dilution, 20 minutes) was performed following incubation in secondary antibodies. Slides were covered using VECTASHIELD mounting medium with DAPI (Vector Laboratories).

Rabbit anti-pS6 (1:10,000; Thermofisher, 44-923G, Lot #2066361) was used in this study. Secondary antibodies were generated in donkey and conjugated to FITC, Cy3 or Cy5 (FITC and Cy5: 1:500, Cy3: 1:1,000; Jackson Immunoresearch Laboratories).

Brain regions were identified based on NeuroTrace labeling by reference to a published betta neuroanatomy atlas^57,58^. pS6+ neurons in each region were counted manually using the Cell Counter feature in FIJI2^59^. Cell counts were normalized by the area of the ROI measured using the Area function in FIJI2.

#### Surgical platform design

The survival surgery platform was adapted from a 3D-printable mouse stereotaxic platform^60^ using Blender software (SFile 3,4). Modifications included resizing the platform and adjusting specific features. A custom holding well was designed to securely accommodate the curved underside of the fish. An additional supporting holder was added to stabilize the body’s sides. Head bars made of disposable transfer pipette tips were secured on either side of the betta’s head. The platform was angled to align the skull of the fish parallel to the stereotaxic platform for surgery.

#### Rostral and caudal Dm lesion surgeries

Prior to surgery, a pulled capillary needle (3.5 inches, Drummond # 3-000-203-G/X; Heat:480, Pull:30, Vel.:80, Time:150, Pressure:200) was painted with DiI (Invitrogen, D3911, 2–3 crystals dissolved in 1 mL ethanol), attached to a vacuum, and mounted to a stereotaxic arm. Fish were then anesthetized by submersion in Tricaine Methanesulfonate stock solution (MS-222; Syndel, 0.08%, pH 7–7.5 using Tris buffer pH 10) diluted 1:10 in water optimized for betta husbandry^51^. Fish were intubated by inserting the tip of a transfer pipette attached to tubing into the mouth to flow tricaine over the gills during surgery. Anesthesia during surgery was a 1:50 dilution of MS-222 stock in betta water and flowed at a rate of 1 mL/minute. For brain surgeries, scales and cutaneous epithelium were then removed over the surgical area using tissue stainless-steel forceps (Dumont) and a cranial window was made using micro dissecting spring scissors (Roboz Surgical Instrument) and forceps.

A cranial window was made in the skull of all fish, whether lesioned or sham-lesioned. The cranial windows extended from the point of calibration (the midpoint of the intersection of the forebrain and the optic lobes) to the target lesion area. Since the rostral Dm is situated farther from the point of calibration compared to caudal Dm, the cranial window for rostral Dm lesions (∼1 mm) was approximately twice the size of the window for caudal Dm lesions (∼0.5 mm). This discrepancy in window size might have contributed to the lower rates of flaring observed in sham-rDm lesioned animals compared to sham-cDm lesioned animal.

Once the cranial window was formed, the DiI covered needle was zeroed at the calibration site. For cDm lesions, the needle was then shifted 200 µm anterior, 0 µm lateral, and 100 µm ventral to the zero point before aspiration. For rDm lesions, the needle was shifted 800 µm anterior, 0 µm lateral, and 100 µm ventral to the zero point. Since Dm is positioned in the medial most part of the forebrain, we performed bilateral lesions by aspirating at the midline with a needle of ∼300 µm diameter. Aspirations were performed by guiding on the vacuum connected to aspiration needle while watching the tissue underneath a stereoscope to ensure proper removal. Following surgery, the small cranial window was sealed using a drop of kwik-sil (World Precision Instruments) and allowed to cure for 6–8 minutes. Fish were returned to their home tanks and allowed to recover for 4 days before post-surgery testing. Following post-surgery behavior, fish were euthanized and tissue was processed to determine the accuracy of the lesion.

#### Tracer Injection

Fish were anesthetized using the method described above. To label retinal ganglion cells, cholera toxin b was injected into the vitreous fluid of the eye through mouth pipetting (∼10 nL), by inserting a capillary needle attached to an aspirator tube through a small hole made with spring scissors. Ctb was allowed to travel for 72 hours before fish were euthanized and tissue processed.

---

**Supplemental Figure 1:**
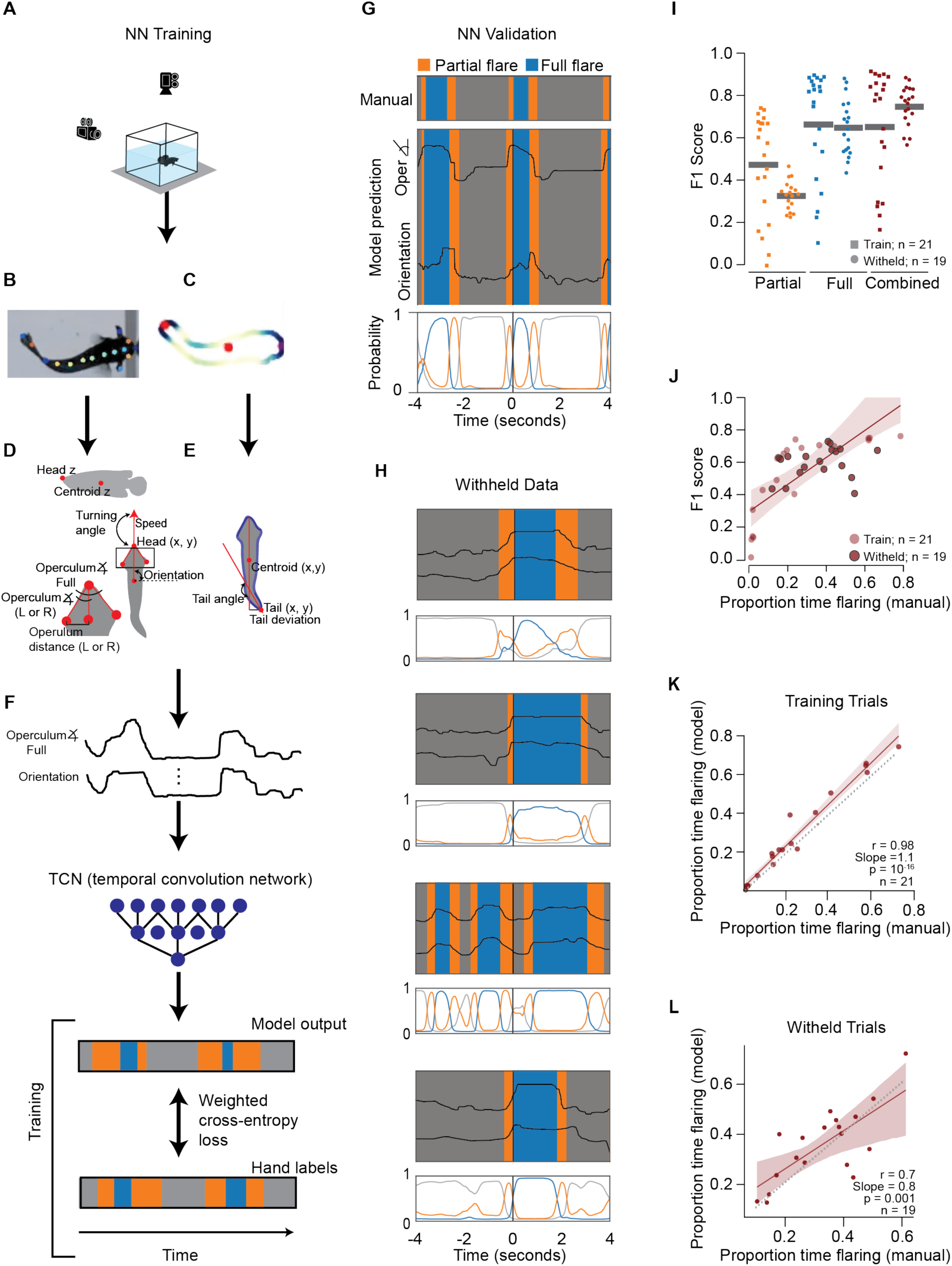
*daart* pipeline for automated scoring of flaring bouts. **(A)** Video setup of betta recorded from the top and side. **(B)** Fish body parts tracked using markerless pose estimation. **(C)** Contour of the body tracked. **(D, E)** Features of shape and motion extracted using **(D)** tracked body parts or **(E)** contour. **(F)** Temporal convolution network (TCN) model training pipeline. **(G)** Comparison of manually-scored flaring events (top), model-scored flaring events (middle), and probability of fish being in each state (background, full, partial flare) according to TCN (bottom). **(H)** Examples of model-scored flaring events in withheld trials. **(I)** F1 scores of training and withheld trials for 3 states (partial flaring, full flaring, and combined (combination of partial and full flaring)). **(J)** Correlation of F1 score and proportion time flaring scored manually. **(K, L)** Correlation of proportion time flaring scored manually or by model for training trials **(K)** and withheld trials **(L)**. **(I, J, K, L)** each dot denotes a trial.

**Supplemental Figure 2:**
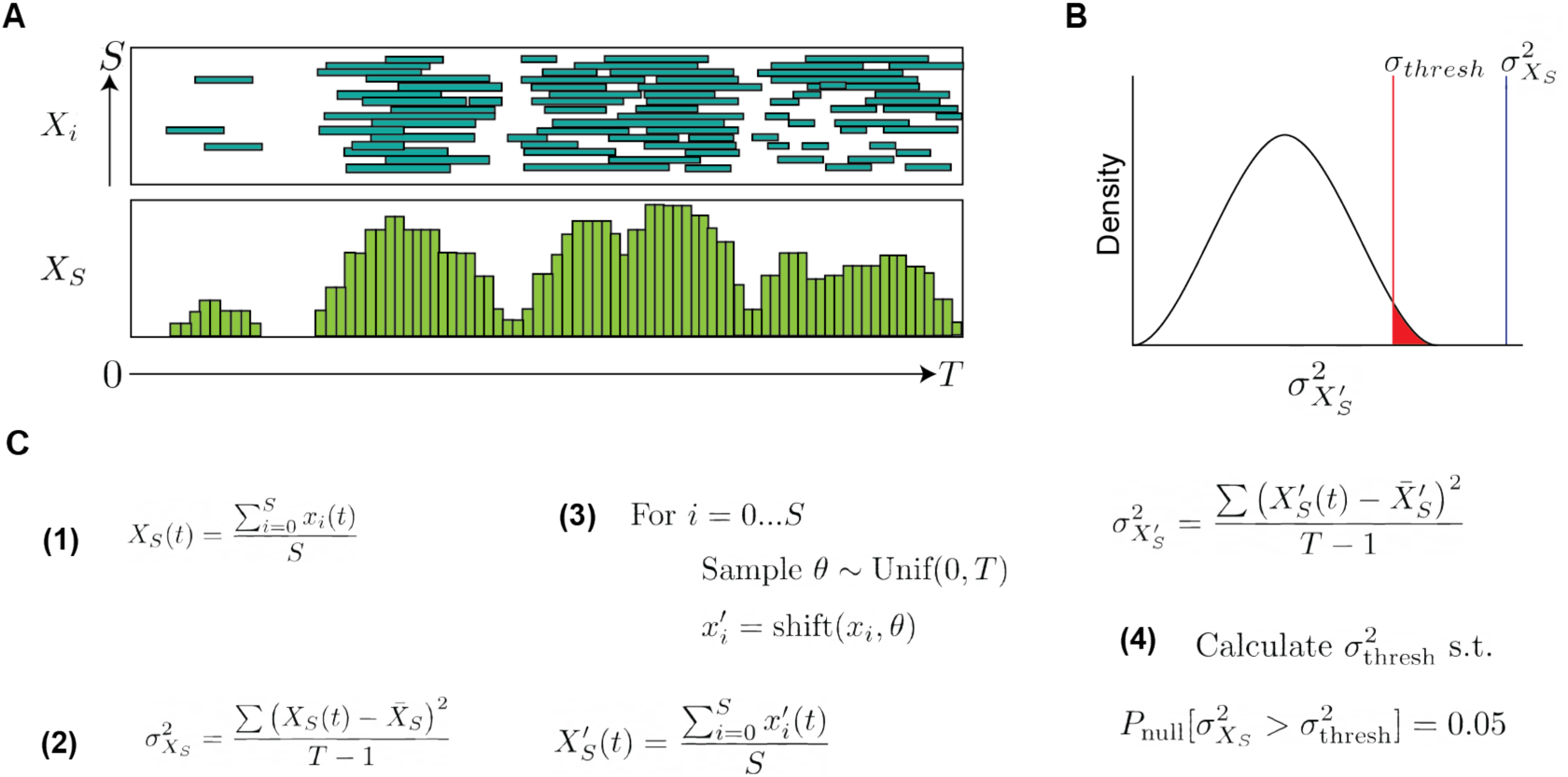
Quantification of synchronization with animations. **(A)** Flare responses (Xi) in each loop of the animation (S) (top) and the flaring frequency (Xs) towards each timepoint in the animation (t) (bottom)) **(B)** Distribution of variance of flaring frequency by chance 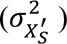 with actual variance of flaring frequency 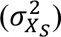 falling significantly outside the distribution by chance, denoting synchronization. **(C)** Equations calculating: the flaring frequency at time (t) (1), the variance of flaring frequencies (2), a set of variances of flaring frequency due to chance (3), whether the actual variance of flaring frequencies falls significantly outside of variance due to chance (4).

**Supplemental Figure 3:**
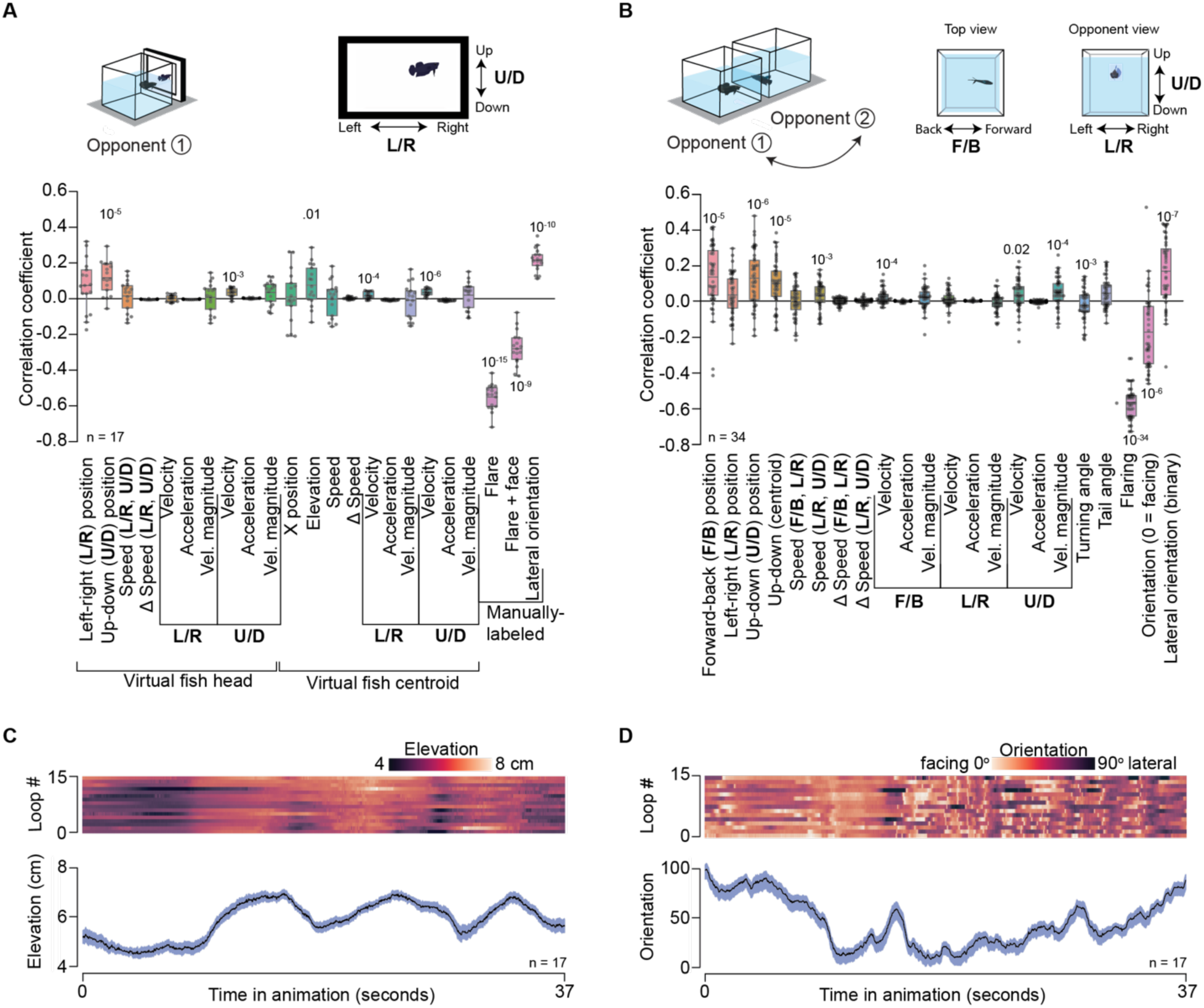
Betta coordinate their flare response, orientation, and elevation to multiple dynamic visual cues emitted by naturalistic animations and conspecifics. (A, B) Correlation between flaring of opponent 1 with behaviors of the animation (A) or opponent 2 (B). p-values by comparing correlations to shuffled controls. (C) Elevation of an individual exposed to animation (top) and population average (bottom). (D) Orientation of an individual exposed to animation (top) and population average (bottom).

**Supplemental Figure 4:**
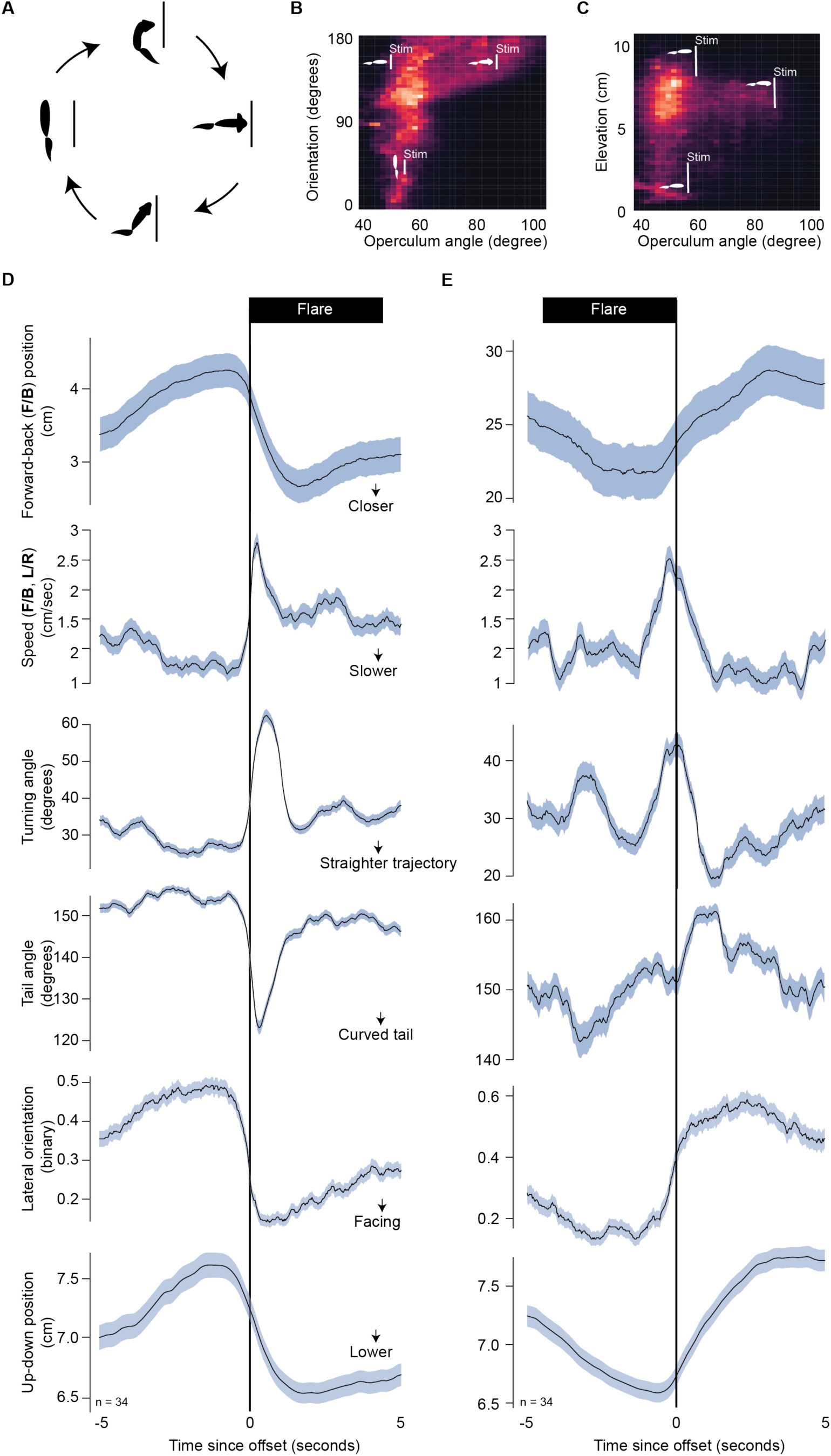
Concurrent dynamics of multiple visual features emitted by betta as they begin or terminate flaring. **(A)** Betta transition between “phases”. Phase starting at the left and moving clockwise: (1) betta lateral and not flaring, (2) betta beginning to extend gills and turn to face, (3) betta flaring and facing, (4) betta beginning to retract gills and turning laterally. **(B, C)** 2D histogram of fish’s operculum angle and orientation **(B)** or elevation **(C)**. White overlayed diagrams denote betta posture. **(D, E)** Peri-event time histograms (mean±SEM) of changes in within-fish dynamics at the onset (left) or offset (right) of flaring bout.

**Supplemental Figure 5:**
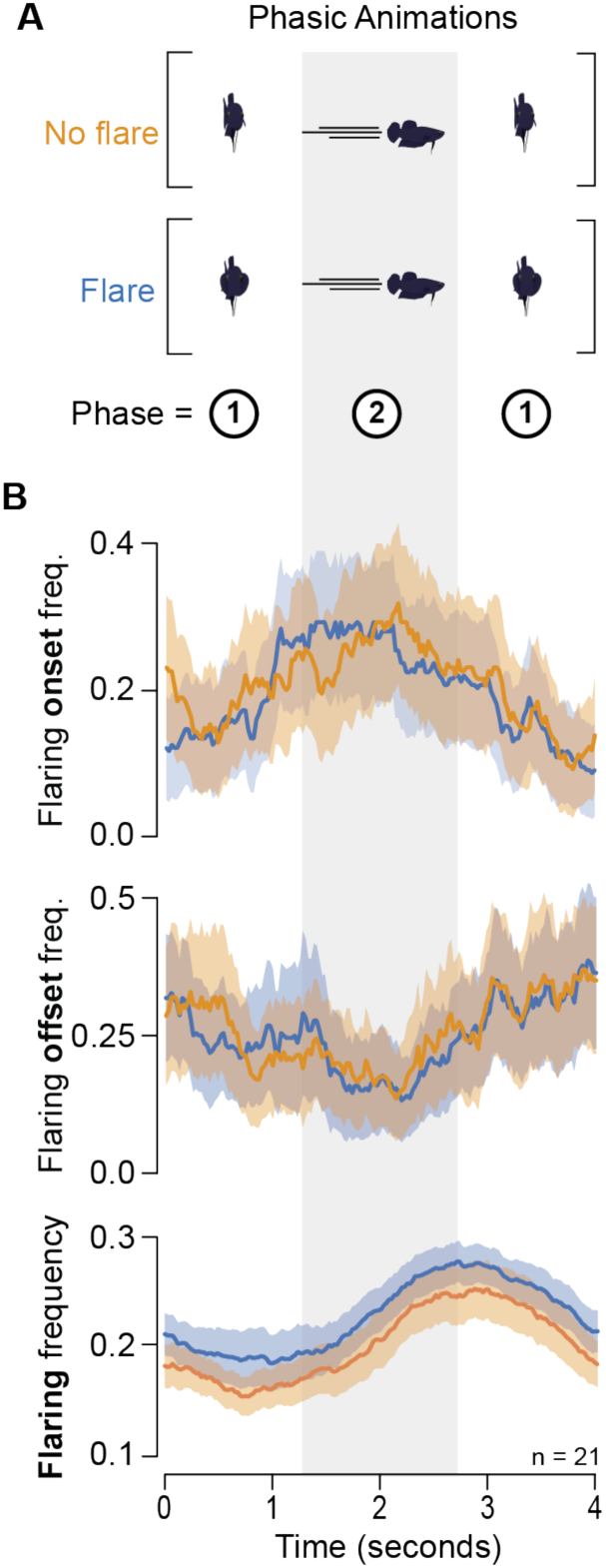
Betta coordinate their flaring with a biphasic animation. **(A)** Animations were composed of two alternating phases with distinct combination of flare, orientation, and speed state. **(B)** (top) Frequency of flaring bout onsets (top) offsets (middle) and overall flaring frequency (bottom) aligned to a loop of the animation.

**Supplemental Figure 6:**
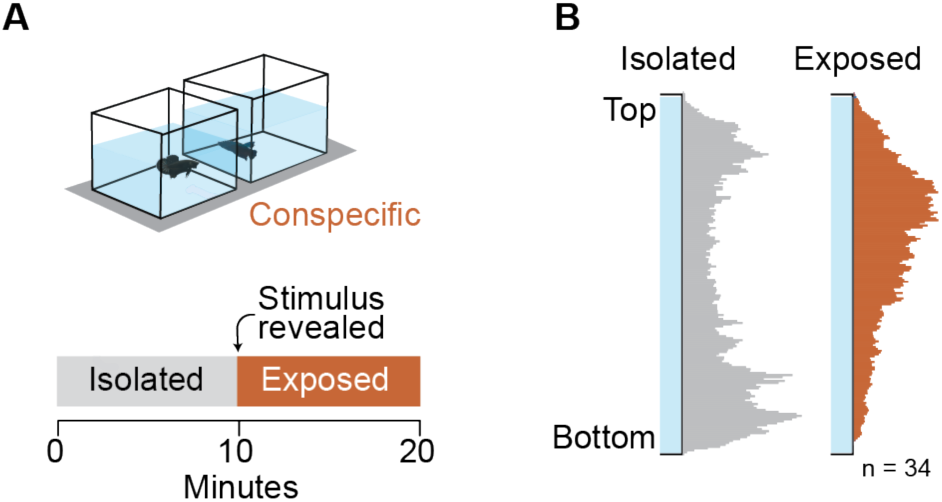
Betta elevate themselves in the water column during display. **(A)** Conspecific paradigm: two opponents in neighboring tanks. **(B)** Elevation distribution during isolation (left) and exposure (right) to opponent.

**Supplemental Figure 7:**
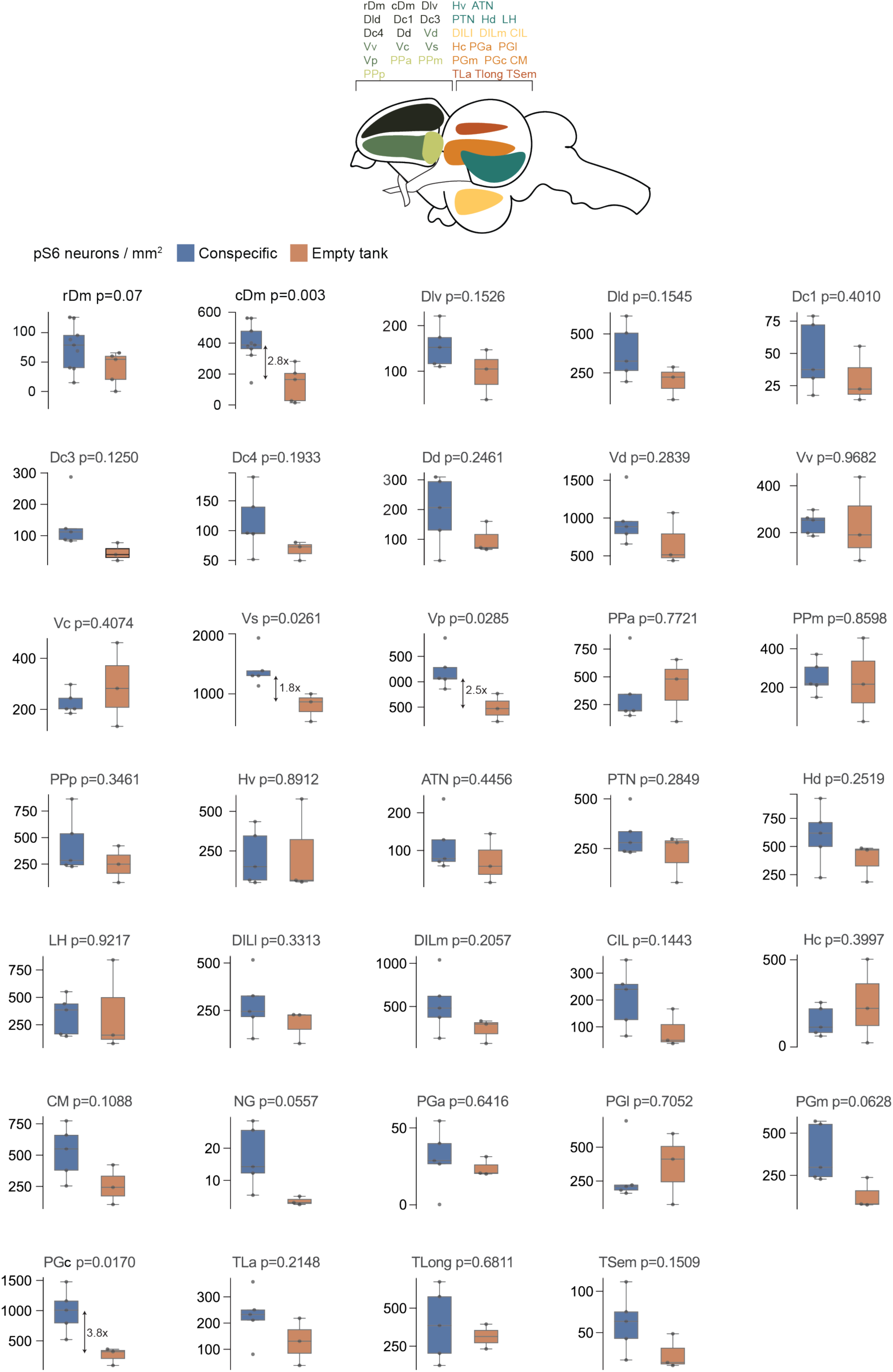
pS6 activity map of areas differentially activated following aggressive display. Dm: dorsal medial area of the pallium, Dlv: ventral division of the lateral zone of the dorsal telencephalon, Dld: dorsal division of the lateral zone of the dorsal telencephalon, Dc: central zone of the dorsal telencephalon, Dd: dorsal zone of the dorsal telencephalon, Vd: dorsal nucleus of the ventral telencephalon, Vv: ventral nucleus of the ventral telencephalon, Vc: central nucleus of the ventral telencephalon, Vs: supracommissural nucleus of the ventral telencephalon, Vp: postcommissural nucleus of the ventral telencephalon, PPa: anterior parvocellular preoptic nucleus, PPm: magnocellular preoptic nucleus, PPp: posterior parvocellular preoptic nucleus, HV: ventral zone of periventricular hypothalamus, ATN: anterior tuberal nucleus, PTN: posterior tuberal nucleus, Hd: dorsal zone of periventricular hypothalamus, LH: lateral hypothalamic nucleus, DILl: lateral diffuse nucleus of the inferior lobe, DILm: medial diffuse nucleus of the inferior lobe, CIL: central diffuse nucleus of the inferior lobe, Hc: caudal zone of periventricular hypothalamus, CM: corpus mammilare (mamillary body), PGa: anterior preglomerular nucleus, PGl: lateral preglomerular nucleus, PGm: medial preglomerular nucleus, PGc: commissural preglomerular nucleus, TLa: anterior torus semicircularis, TLong: torus longitudinalis, TSem: torus semicircularis. p-values by unpaired t-test.

**Supplemental Figure 8:**
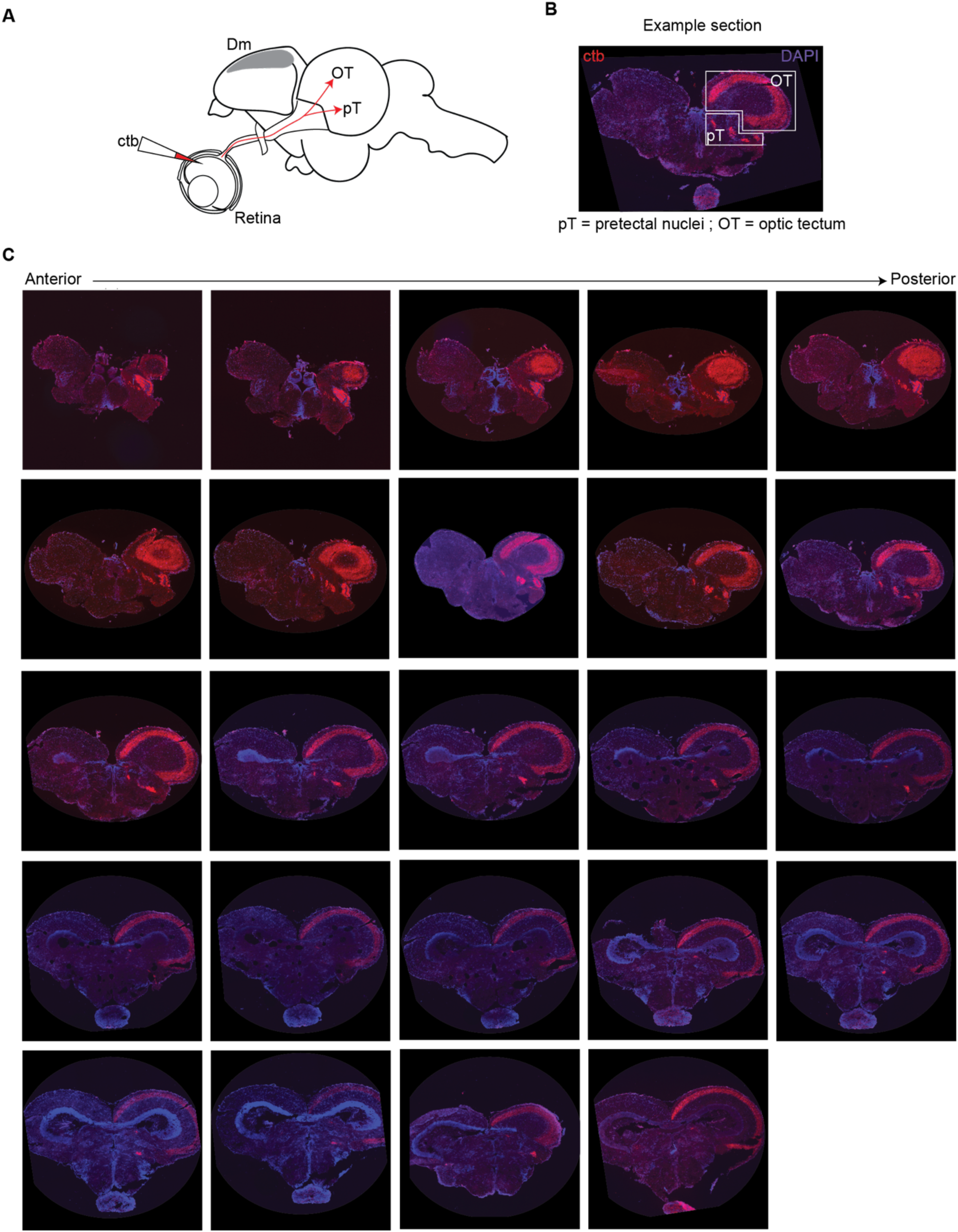
Retinorecipient nuclei in adult betta. **(A)** Schematic diagram of injection site (intraocular) and presumptive retinal projections. **(B)** Labelled retinal ganglion cell (RGC) terminals in the optic tectum (OT) and a group of pretectal nuclei (pT). **(C)** RGC termination in OT and pT along anterior-posterior axis. Naming derived from Vigouroux et al. (ref. ^45^).

**Supplemental Figure 9:**
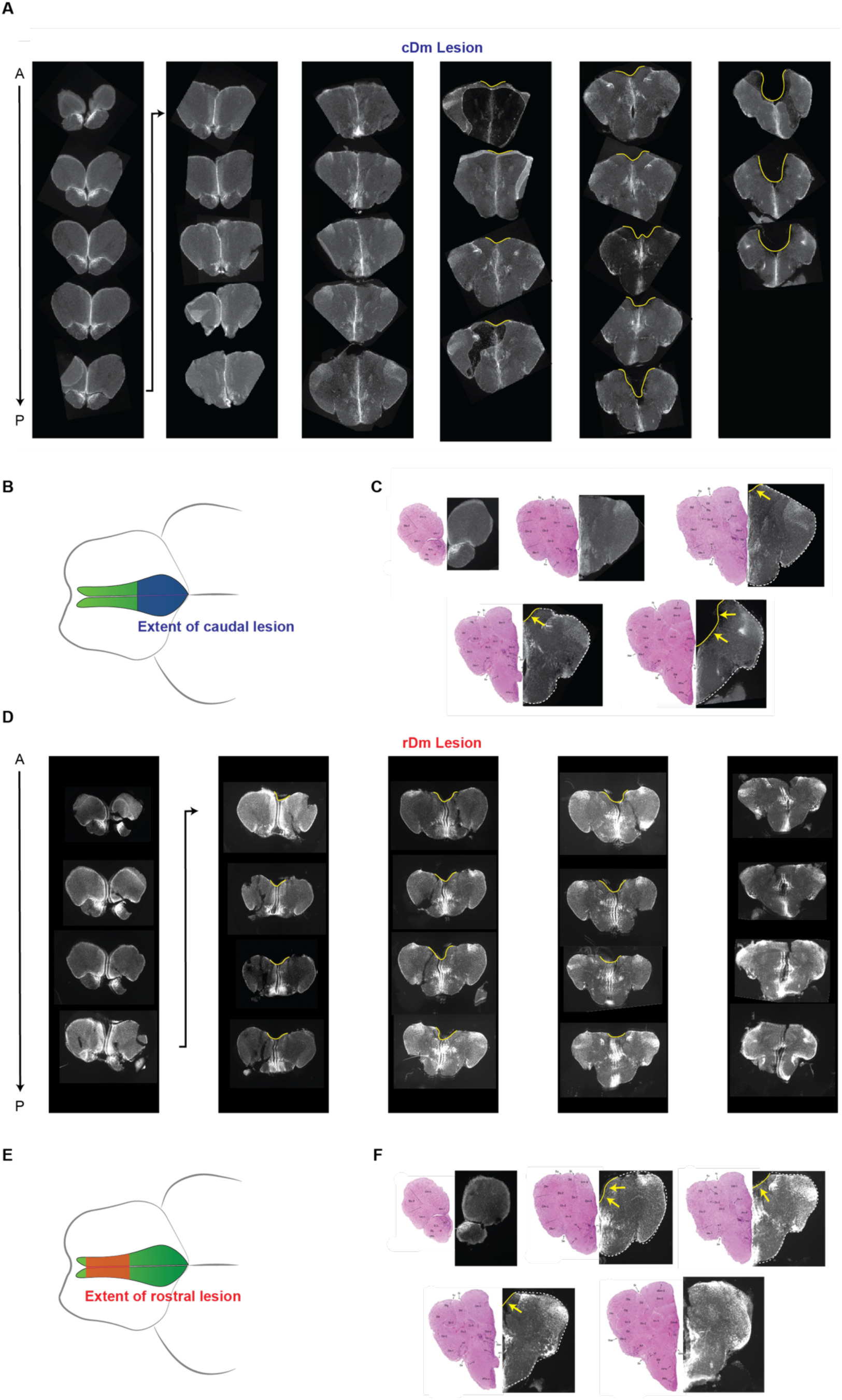
Histology of rostral and caudal Dm lesions. **(A)** Coronal sections of forebrain with caudal lesion. **(B)** Diagram of extent of lesion along anteroposterior axis. **(C)** Select sections from series aligned to published atlas. **(D)** Coronal sections of forebrain with rostral lesion. **(E)** Diagram of extent of lesion along anteroposterior axis. **(F)** Select sections from series aligned to published atlas^57^. Arrows point to the brain region removed by the lesion.

**Supplemental Figure 10:**
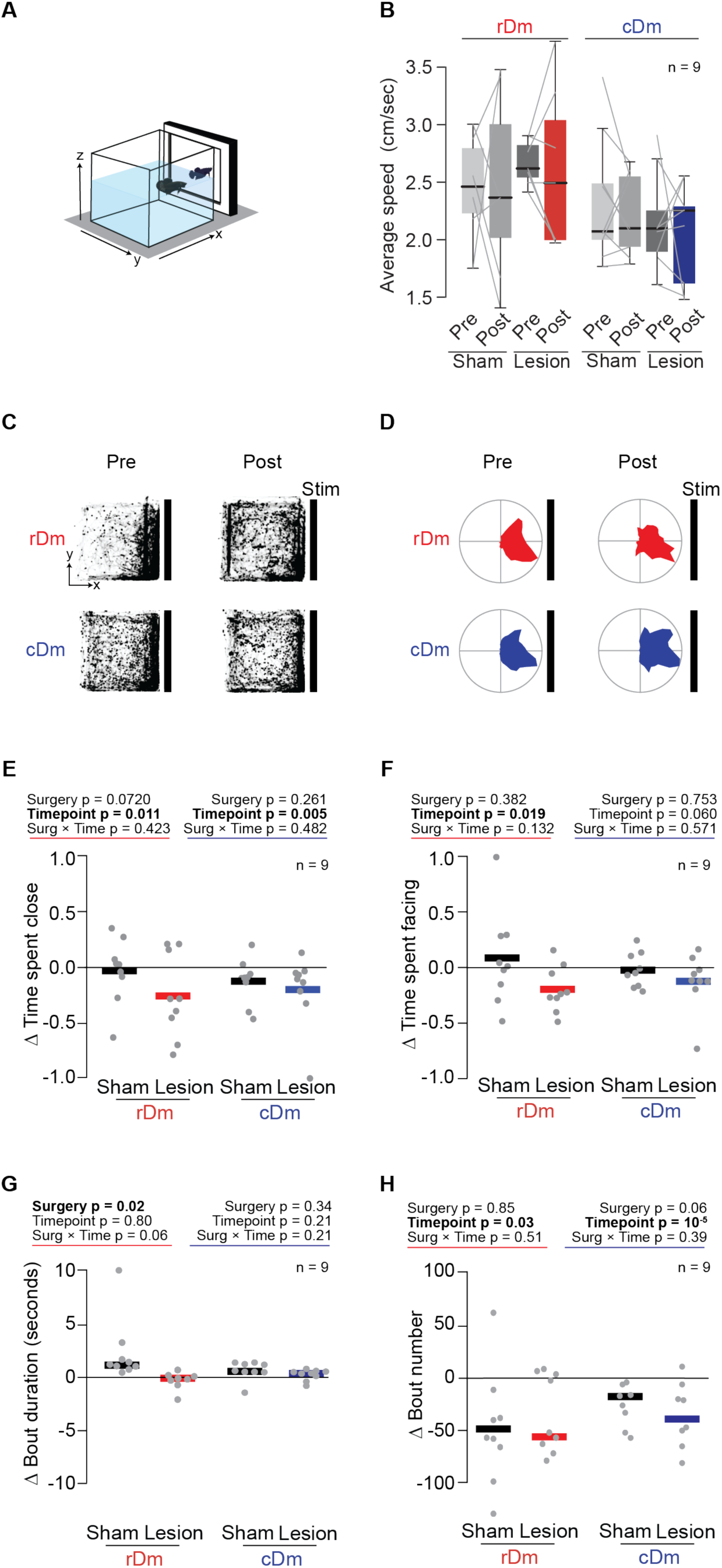
Additional behavioral metrics towards animation following lesions. **(A)** Naturalistic animation paradigm: one opponent faces a naturalistic animation modeled after a male displaying aggression. **(B)** Average speed during free swimming (before the animation starts) pre- and post-caudal Dm (left) and rostral Dm (right) lesion and associated sham controls. **(C)** Heatmaps showing tank occupancy (from a top view) by fish during exposure pre- and post-lesion. **(D)** Polar plots showing distributions of the head orientation during exposure pre- and post-lesion. **(E)** Change in the proportion time spent in the quarter of the tank closest to the stimulus. **(F)** Change (post-pre surgery) in the proportion time spent facing within 180° of the stimulus. **(G)** Change (post-pre surgery) in flare bout duration. **(H)** Change (post-pre surgery) in the number of flare bouts. Dots denote individuals. p-values by mixed models ANOVA.

**Supplemental Figure 11:**
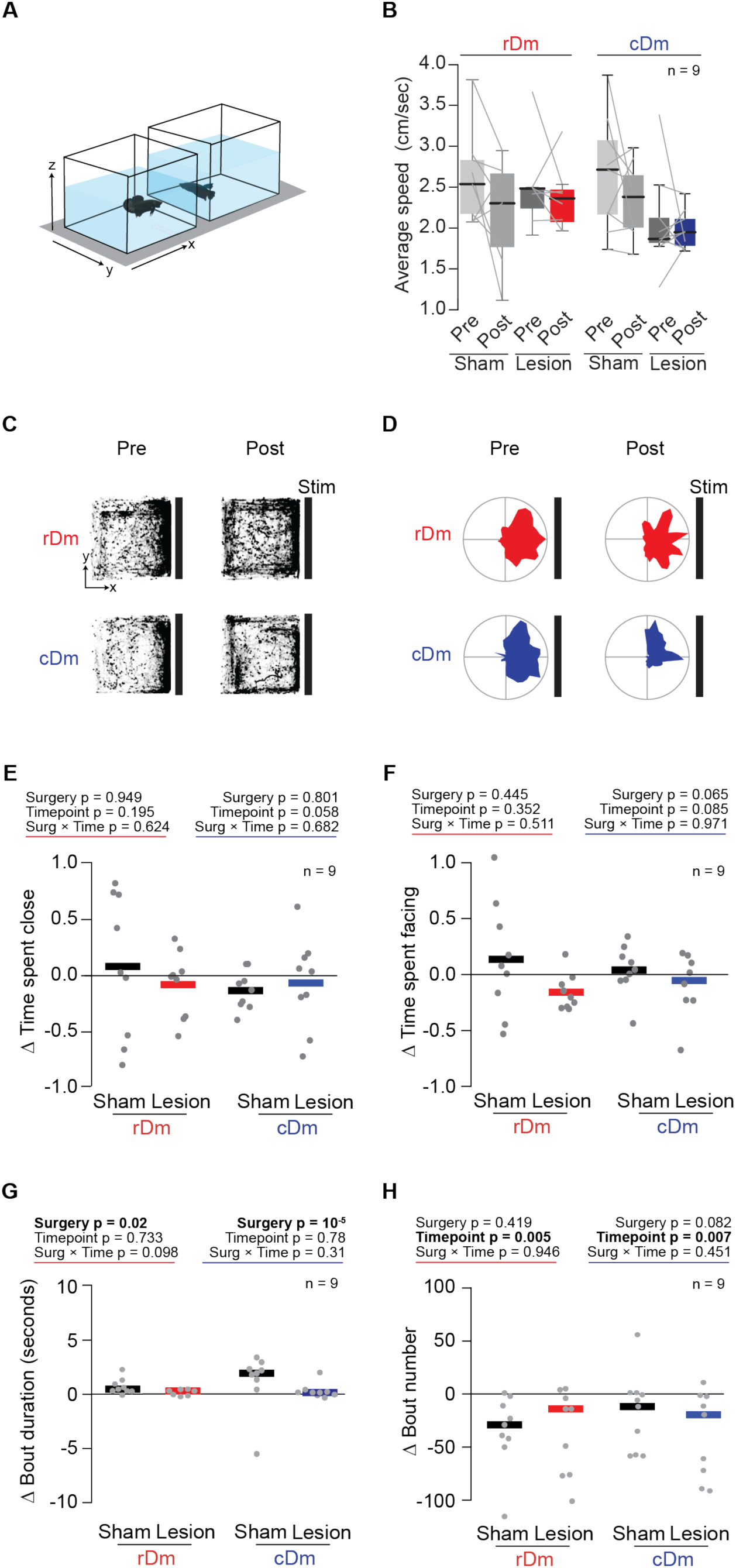
Additional metrics of arousal, attention, and aggression towards conspecific following lesion. **(A)** Conspecific paradigm: two opponents in neighboring tanks. **(B)** Average speed during free swimming (before exposure to conspecific starts) pre- and post-caudal Dm (left) and rostral Dm (right) lesion and associated sham controls. **(C)** Heatmaps showing tank occupancy (from a top view) by fish during exposure pre- and post-lesion. **(D)** Polar plots showing distributions of the head orientation during exposure pre- and post-lesion. **(E)** Change in the proportion time spent in the quarter of the tank closest to the stimulus. **(F)** Change (post-pre surgery) in the proportion time spent facing within 180° of the stimulus. **(G)** Change (post-pre surgery) in flare bout duration. **(H)** Change (post-pre surgery) in the number of flare bouts. Dots denote individuals. p-values by mixed models ANOVA.

**Supplemental Figure 12:**
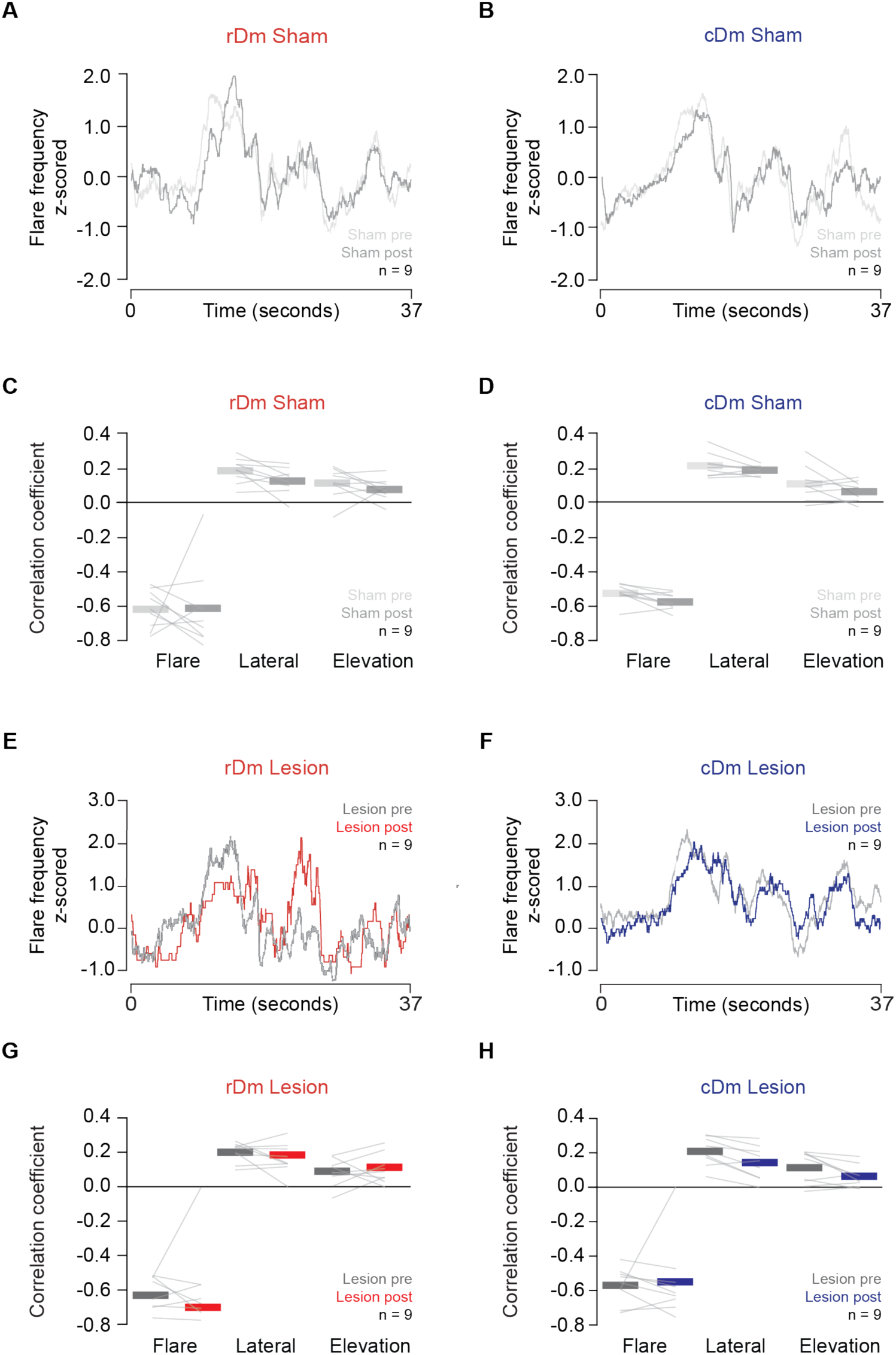
Coordination of flare response following sham and lesion surgeries. **(A, B)** Average flaring frequency towards animation pre- and post-sham surgery associated with rDm **(A)** and cDm **(B)** lesion. **(C, D)** Correlation between flaring of sham-operated animals as controls of rDm **(C)** or cDm **(D)** lesions pre- and post-surgery with behaviors of the animation. **(E,F)** Average flaring frequency towards animation pre- and post-rDm **(E)** and cDm **(F)** lesion. **(G, H)** Correlation between flaring of rDm **(G)** or cDm **(H)** lesioned fish pre- and post-surgery with behaviors of the animation.

