## Supplemental File 2 for "Coordination and persistence of aggressive visual communication in Siamese fighting fish": auto_filter_full.py

```
#!/usr/bin/env python3
# -*- coding: utf-8 -*-
"""
Created on Thu Jun  4 13:06:14 2020

"""

import numpy as np
import pandas as pd

def auto_scoring_tracefilter_full(data,p=0.5,p_tail=15,p_head=5,angle_tol=75):
    mydata = data.copy()
    spine_column=['A_head',"F_spine1",'mid_spine1_spine2',"G_spine2",'mid_spine2_spine3',"H_spine3","I_spine4","J_spine5","K_spine6","L_spine7","B_rightoperculum",
                 'E_leftoperculum']
    for i,c in enumerate(spine_column):
        #likelihood check
        likelihood_check=np.array(data[c]['likelihood']<p)
        mydata.loc[likelihood_check,(c,'x')]=np.nan;  mydata.loc[likelihood_check,(c,'y')]=np.nan
       
        #position difference check

        #head and gill distance check
        if spine_column[i] in ['A_head',"B_rightoperculum",'E_leftoperculum']:
            if c=="A_head":
                #cal dist between head->spine1/operculum->head, if it's too large discard it
                head_spine1=np.array([mydata['A_head']['x']-mydata['F_spine1']['x'],mydata['A_head']['y']-mydata['F_spine1']['y']]).T
                head_dist=np.sqrt(np.sum(head_spine1*head_spine1,axis=1))
                head_dist_check=head_dist>25 #assume normal, use 3 \sigma rule
                mydata.loc[head_dist_check,(c,'x')] = np.nan; mydata.loc[head_dist_check,(c,'y')] = np.nan
         
            else:
                gill_head=np.array([mydata[c]['x']-mydata['A_head']['x'],mydata[c]['y']-mydata['A_head']['y']]).T
                gill_dist=np.sqrt(np.sum(gill_head*gill_head,axis=1))
                gill_dist_check=gill_dist>75
                mydata.loc[gill_dist_check,(c,'x')] = np.nan; mydata.loc[gill_dist_check,(c,'y')] = np.nan
                mydata.loc[gill_dist_check,("A_head",'x')] = np.nan; mydata.loc[gill_dist_check,("A_head",'y')] = np.nan
                
        #angle check
         #use the line from spine1 to mid point of spine1,spine2 as baseline, any segment to fit the spline should not have a angle
            #with the base line larger than 75 degree.
        if spine_column[i]=='mid_spine1_spine2':
            baseline=np.array([mydata['mid_spine1_spine2']['x']-mydata['F_spine1']['x'],mydata['mid_spine1_spine2']['y']-mydata['F_spine1']['y']]).T
        if spine_column[i] in ["G_spine2",'mid_spine2_spine3',"H_spine3","I_spine4","J_spine5","K_spine6","L_spine7"]:
            orientation1=np.array([mydata[spine_column[i]]['x']-mydata[spine_column[i-1]]['x'],mydata[spine_column[i]]['y']-mydata[spine_column[i-1]]['y']]).T
            orientation2=np.array([mydata[spine_column[i]]['x']-mydata[spine_column[i-2]]['x'],mydata[spine_column[i]]['y']-mydata[spine_column[i-2]]['y']]).T
            orientation3=np.array([mydata[spine_column[i]]['x']-mydata[spine_column[i-3]]['x'],mydata[spine_column[i]]['y']-mydata[spine_column[i-3]]['y']]).T
            orientation=np.copy(orientation1); mask=np.isnan(np.sum(orientation,axis=1))           #if the previous point is already na, check the vector to the second closest point
            orientation[mask]=orientation2[mask]; mask=np.isnan(np.sum(orientation,axis=1))     # if it's also na,use next previous point ,and if the second closest point is also nan, stop here                        
            orientation[mask]=orientation3[mask]; mask=np.isnan(orientation) #if the previous 3 points are all NA, just.....don't use it for safety reason.
            safety_check=np.sum(mask,axis=1)!=0
            #if the baseline contains nan, skip this step
            inner_product =np.sum(baseline*orientation,axis=1)
            cos=inner_product/np.sqrt(np.sum(baseline*baseline,axis=1))/np.sqrt(np.sum(orientation*orientation,axis=1))
            angle=np.arccos(cos)/np.pi*180;angle_check=np.logical_or(np.logical_and(np.invert(np.isnan(angle)),angle>angle_tol),safety_check)          
            mydata.loc[angle_check,(c,'x')]=np.nan; mydata.loc[angle_check,(c,'y')]=np.nan           
    orientation=-head_spine1
    inner_product =np.sum(baseline*orientation,axis=1)
    cos=inner_product/np.sqrt(np.sum(baseline*baseline,axis=1))/np.sqrt(np.sum(orientation*orientation,axis=1))
    angle=np.arccos(cos)/np.pi*180;angle_check=np.logical_and(np.invert(np.isnan(angle)),angle>angle_tol)
    mydata.loc[angle_check,('A_head','x')]=np.nan; mydata.loc[angle_check,('A_head','y')]=np.nan

    return mydata

def midpoint_wLikelihood (x1, y1, l1,x2, y2,l2):
    '''
    give definition x1: x-value object 1, y1: y-value object 1, x2: x-value object 2
    y2: y-value object 2, l1: likelihood object 1,l2:likelihood object 2
    '''
    midpointx = (x1 + x2)/2
    midpointy = (y1 + y2)/2
    MinLikelihood=np.minimum(l1,l2) #the likelihood is set to the minimum of 2 columns

    return list(zip(midpointx, midpointy,MinLikelihood))

def transform_data(df):
    mid_spine1_spine2=midpoint_wLikelihood(df['F_spine1']['x'],df['F_spine1']['y'],df['F_spine1']['likelihood'],
                                           df['G_spine2']['x'],df['G_spine2']['y'],df['G_spine2']['likelihood'])
    name_arr=[["mid_spine1_spine2","mid_spine1_spine2","mid_spine1_spine2"], ['x', 'y', 'likelihood']]
    mid_spine1_spine2=pd.DataFrame(mid_spine1_spine2,columns=pd.MultiIndex.from_arrays(name_arr,names=["bodyparts",'coords']))
    df.reset_index(drop=True, inplace=True)
    df=pd.concat([df,mid_spine1_spine2],axis=1)
    mid_spine2_spine3=midpoint_wLikelihood(df['G_spine2']['x'],df['G_spine2']['y'],df['G_spine2']['likelihood'],
                                           df['H_spine3']['x'],df['H_spine3']['y'],df['H_spine3']['likelihood'])
    name_arr=[["mid_spine2_spine3","mid_spine2_spine3","mid_spine2_spine3"], ['x', 'y', 'likelihood']]
    mid_spine2_spine3=pd.DataFrame(mid_spine2_spine3,columns=pd.MultiIndex.from_arrays(name_arr,names=["bodyparts",'coords']))
    df=pd.concat([df,mid_spine2_spine3],axis=1)
    return df

def filter_tailbeating(data,p0=50,p_head = 30, p1=25, p2 = 10, t1 = 20):
    # check points location intervals
    mydata = data.copy()
    spine_column=["A_head","F_spine1","G_spine2","H_spine3","I_spine4","J_spine5","K_spine6","L_spine7"] #,'D_tailtip','C_tailbase']
    for i,c in enumerate(spine_column):
        # using the spine1 as the original points
        if i == 0:
            dist = np.sqrt(np.square(data[spine_column[i+1]]['x']-data[c]['x'])+np.square(data[spine_column[i+1]]['y']-data[c]['y']))
            dist_check = dist > p_head
            mydata[c]["x"][dist_check]  = np.nan
            mydata[c]["y"][dist_check]  = np.nan
        if (i>1 and i<(len(spine_column)-1)):
            r_decision = False
            dist1=np.sqrt(np.square(data[spine_column[i-1]]['x']-data[c]['x'])+np.square(data[spine_column[i-1]]['y']-data[c]['y']))
            dist2=np.sqrt(np.square(data[spine_column[i+1]]['x']-data[c]['x'])+np.square(data[spine_column[i+1]]['y']-data[c]['y']))
            # further check the relative position:
            if i > 2:
                dist3 = np.sqrt(np.square(data["F_spine1"]['x']-data[c]['x'])+np.square(data["F_spine1"]['y']-data[c]['y']))
                if np.logical_or((dist3[0] > ((i-1)*p1+t1)),(dist3[0]<((i-3)*p2))):
                    r_decision = True
            dist_check= np.logical_or(((dist1>p1)|(dist2>p1)), r_decision)
            mydata[c]["x"][dist_check] = np.nan
            mydata[c]["y"][dist_check] = np.nan
        if i==(len(spine_column)-1):
            dist1=np.sqrt(np.square(data[spine_column[i-1]]['x']-data[c]['x'])+np.square(data[spine_column[i-1]]['y']-data[c]['y']))
            dist2=np.sqrt(np.square(data[spine_column[i-2]]['x']-data[c]['x'])+np.square(data[spine_column[i-2]]['y']-data[c]['y']))
            dist3 = np.sqrt(np.square(data["F_spine1"]['x']-data[c]['x'])+np.square(data["F_spine1"]['y']-data[c]['y']))
            r_decision = np.logical_or(dist3[0]>((i-1)*p1), dist3[0]<((i-4)*p2))
            dist_check=np.logical_or(((dist1>p1)|(dist2>p1)), r_decision)
            mydata[c]["x"][dist_check] = np.nan
            mydata[c]["y"][dist_check] = np.nan
        
    return mydata
```
