## Supplemental File 2 for "Coordination and persistence of aggressive visual communication in Siamese fighting fish": contour_auto_single_FINALT.py

```
#!/usr/bin/env python3
# -*- coding: utf-8 -*-
"""
Created on Wed Mar 16 10:45:48 2022

"""
import sys
import numpy as np
import pandas as pd
import cv2
from tqdm import tqdm
from scipy.ndimage import zoom  
from contour_utils import *
from find_features import features
from scipy import sparse
from LoopingArray import LoopingArray
import pickle
import matplotlib.pyplot as plt
import os
from glob import glob


def find_background(video_path,sample_length = None,image_shape = (500,500,3)):
    '''
    sample_length: number of samples used to calculate background, default to full video
    '''
    if not os.path.exists(video_path):
        raise Exception("video path not exist")
    cap = cv2.VideoCapture(video_path) 
    frames = np.zeros(image_shape) #running mean of video
    if sample_length is None:
        sample_length = float("inf")
    n_sample = 0
    while True:
        ret, frame = cap.read()
        n_sample += 1
        if not ret or n_sample>sample_length:
            break
        frame = cv2.cvtColor(frame,cv2.COLOR_BGR2RGB)
        frames = frames*(n_sample-1)/n_sample + 1/n_sample*frame 
        if n_sample % 5000 == 0:
            print("{} samples processed".format(n_sample))
    print("background for {}".format(video_path))
    plt.figure()
    cv2.imwrite(os.path.join(output_dir, 'background.png'), frames)
    return frames

def pre_filter(background, frame):
    # get background mask
    background = cv2.imread(background)
    imgray_back = cv2.cvtColor(background, cv2.COLOR_BGR2GRAY)
    ret, thresh_back = cv2.threshold(imgray_back, 50, 255, 0)
    thresh_back_flip = abs(thresh_back.astype(float) - 255)
    # get the frame mask
    imgray_fore = cv2.cvtColor(frame, cv2.COLOR_BGR2GRAY)
    ret, thresh_fore = cv2.threshold(imgray_fore, 50, 255, 0)
    thresh_fore_flip = abs(thresh_fore.astype(float) - 255)

    subtraction  = cv2.subtract(thresh_fore_flip, thresh_back_flip)

    cv2.imwrite(os.path.join(output_dir, 'subtraction.png'), subtraction)
    
    im = cv2.imread(os.path.join(output_dir, 'subtraction.png'))
    return im

def find_contour(videopath,head,length=40*300,start=0,step=1,pre_filter=None,threshold = 50,shrink_kernel = 3):
    '''
    prefilter:a function that changes mask array
    length: Total number of frames needed
    start: which frame to start
    step: pick every * frame
    thres: thresholding parameter when doing cv2.threshold
    shrink_kernel: how much you want to shrink the thresholded mask to avoid reflections
    '''
    background = find_background(videopath, sample_length = 50000, image_shape = (500, 500, 3))
    vidcap = cv2.VideoCapture(videopath)
    contour_array=[]
    length=int(length/step)
    index=start
    vidcap.set(1,index)
    first = True
    for i in tqdm(range(length)):
        success,image_first=vidcap.read()
        if success!=1:
            print("process stops early at {}th iteration".format(i))
            break
        if pre_filter is not None:            
            image = np.invert(pre_filter(os.path.join(output_dir, 'background.png'), image_first))
        index+=step
        vidcap.set(1,index)
        gray = cv2.cvtColor(image, cv2.COLOR_BGR2GRAY)
        ret,th = cv2.threshold(gray, threshold, 255, cv2.THRESH_BINARY)

        if shrink_kernel>0:
          k2 = cv2.getStructuringElement(cv2.MORPH_RECT, (shrink_kernel,shrink_kernel))
          th = cv2.dilate(th, k2, iterations=1)

        contours, hierarchy = cv2.findContours(np.invert(th.copy()), cv2.RETR_TREE, cv2.CHAIN_APPROX_SIMPLE)#simplified contour pts
        #finding the contour with largest area
        fish_contour,flag=find_largest_contour(contours)
        if flag==0:
            print("no valid fish contour find at index {}".format(i))
            img=np.float32(np.full(image.shape,0))#just in case there's no valid contour, won't happen in the current case
        else:
            #draw only the mask of this contour
            img=cv2.drawContours(np.float32(np.full(image.shape,255)),[fish_contour],0,(0,0,0),cv2.FILLED)
        img=cv2.cvtColor(img,cv2.COLOR_BGR2GRAY)

        if shrink_kernel>0:
          k3=cv2.getStructuringElement(cv2.MORPH_RECT, (shrink_kernel+2,shrink_kernel+2))
        else:
          k3=cv2.getStructuringElement(cv2.MORPH_RECT, (3,3))
        img=np.invert(np.array(cv2.erode(img,k3,iterations=1),np.uint8))
        #smoothing
        img=cv2.medianBlur(img, 5)
        true_contour, hierarchy = cv2.findContours(img.copy(), cv2.RETR_TREE, cv2.CHAIN_APPROX_NONE)#complete contour pts
        true_contour,flag=find_largest_contour(true_contour)
        if flag == 0:
          contour_array.append(np.random.randn(100,1,2))
          continue
        conservative_mask = np.array(cv2.drawContours(np.float32(np.full((img.shape[0],img.shape[1]),255)),[true_contour],0,(0,0,0),cv2.FILLED),np.uint8)
        conservative_contour = true_contour
        #re-thresholding image and try to include tail into the contour
        ret,th = cv2.threshold(gray, threshold, 255, cv2.THRESH_BINARY)
        k1 = cv2.getStructuringElement(cv2.MORPH_RECT, (5,5))
        th = cv2.erode(th, k1, iterations=1)

        xbar,ybar=find_centroid(conservative_contour)
        dists=compute_dist(conservative_contour,xbar,ybar)
        thres=np.quantile(dists,0.4)
        y,x=np.nonzero(np.invert(th))#black area, i.e. fish body
        head_x=head.x.iloc[i]
        head_y=head.y.iloc[i]
        th=exclusion(y,x,thres,xbar,ybar,head_x,head_y)           
        th=np.minimum(th,conservative_mask)
        th=np.invert(th)
        contours, hierarchy = cv2.findContours(th, cv2.RETR_TREE, cv2.CHAIN_APPROX_SIMPLE)
        fish_contour,flag=find_largest_contour(contours)
        if flag==0:
            print("no valid fish contour find at index {}".format(index-1))
            img=np.float32(np.full(image.shape,0))#just in case there's no valid contour, won't happen in the current case
        else:
            img=cv2.drawContours(np.float32(np.full(image.shape,255)),[fish_contour],0,(0,0,0),cv2.FILLED)
        img=cv2.cvtColor(img,cv2.COLOR_BGR2GRAY)
        #to get rid of small holes in the mask
        k3=cv2.getStructuringElement(cv2.MORPH_RECT, (3,3))
        img=np.invert(np.array(cv2.erode(img,k3,iterations=1),np.uint8))
        img=cv2.medianBlur(img, 5)
        true_contour, hierarchy = cv2.findContours(img.copy(), cv2.RETR_TREE, cv2.CHAIN_APPROX_NONE)#complete contour pts
        true_contour,_=find_largest_contour(true_contour,interpolate=1)
        try:
            contour_array.append(true_contour)
        except:
            contour_array.append(np.zeros((1,1,2),dtype=np.float64))
        
        if first: #plot sample contour 
            sample_image = cv2.drawContours(image_first.copy(),[true_contour],0,(255,0,0),2)
            plt.savefig("sample_contour.png")
            print("sample contour of the first frame has been saved to sample_contour.png")
            first = False
        
    return contour_array

def exclusion(y,x,thres,xbar,ybar,head_x,head_y):
    dists1=compute_dist(np.vstack([x,y]).T,xbar,ybar)
    #new found part should not be too close to the centroid(as it is tail)
    flag1=dists1<thres
    dists2=compute_dist(np.vstack([x,y]).T,head_x,head_y)
    #new found part shall not be too close to the head, otherwise it is likely a reflection
    flag2=dists2<thres
    flag=np.logical_or(flag1,flag2)
    x=x[~flag]
    y=y[~flag]
    data=np.full((len(x),),255)
    output=np.invert(np.uint8(sparse.coo_matrix((data,(y,x)),shape=(IMAGELENGTH,IMAGELENGTH)).toarray()))
    return output 

def head_on_contour(head_x,head_y,contour):
    dists=compute_dist(contour,head_x,head_y)
    index=np.argmin(dists)
    return index

def compute_cos_fullbody(contour, step=30):
    #compute the curvature for full body
    curviness=np.zeros((0,3),dtype=np.float64)
    contour=contour.squeeze()
    N = len(contour)
    def find_next(i,step):
        return contour[(i+step) % N]
    def find_prev(i,step):
        return contour[(i -step) % N]
    for i in range(N):
        x_cur, y_cur = contour[i]
        x_next, y_next = find_next(i,step)
        x_prev, y_prev = find_prev(i,step)
        vec1=np.array([x_next-x_cur,y_next-y_cur])
        vec2=np.array([x_prev-x_cur,y_prev-y_cur])
        cos=np.sum(vec1*vec2)/np.sqrt(np.sum(vec1*vec1)*np.sum(vec2*vec2))
        curviness=np.concatenate((curviness,np.array([x_cur,y_cur,cos+1]).reshape(1,3)))
    return curviness

def averageBlur(arr,neighbor_width=30):
    #updating an array of curviness
    out=arr.copy()
    l=len(arr)
    for i in range(l):
        out[i]=np.sum(LoopingArray(arr)[(i-neighbor_width):i+neighbor_width+1])/(2*neighbor_width+1)
    return out

def combine_small_segment(value,length,minimal_length=70):
    l=len(value)
    flag=0
    #check if the starting point is a flipping point
    if value[0]==value[l-1]:
        temp=length[l-1]
        flag=1
        length[0]+=temp
        value=value[:l-1]
        length=length[:l-1]
    new_val=[]
    new_length=[]
    stack=0
    for i in range(len(value)):
        ll=length[i]
        #if the segment is smaller than certain length, put it to a existing segment
        #if there is no such thing, prepare that for a future valid segment
        if ll<minimal_length:
            if new_val:                
                new_length[len(new_length)-1]+=ll
            else:
                stack+=ll
        else:
            if len(new_val)==0 or new_val[::-1][0]!=value[i]:
                new_val.append(value[i])
                new_length.append(length[i]+stack)
                stack=0
            else:
                new_length[len(new_length)-1]+=ll
    if flag==1:
        #put the length back when we first cut off the last piece at first
        if new_val[len(new_length)-1]!=new_val[0]:
            new_val.append(new_val[0])
            new_length.append(temp)
        else:
            new_length[len(new_length)-1]+=temp
        new_length[0]-=temp
            
    return np.array(new_val),np.array(new_length)

def predict_tail(contour,head_index,step=None,quantile=0.4,neighbor_width=None,minimal_seg_length=23):
    #default value, in case the image is/is not zoomed
    if step is None:
        step=[int(len(contour)/10),int(len(contour)/10)*1.5]
    if neighbor_width is None:
        neighbor_width=[int(len(contour)/20),int(len(contour)/20)]
    xbar,ybar=find_centroid(contour)
    dists=compute_dist(contour,xbar,ybar)
    thres=np.quantile(dists,quantile)
    #get whether the pts are far from the centroid
    validity=dists>thres
    l=len(dists)
    value,length=runLengthEncoding(validity)
    value,length=combine_small_segment(value, length,minimal_seg_length)
    def find_segment(value,length):
        #index of segment
        l=len(value)
        new_length=length.copy()
        new_value=value.copy()
        #append the last segment unfinished segment to the front to compare the length
        if value[l-1]==value[0]:
            new_length[0]=length[0]+length[l-1]
            new_value=value[:-1]
            new_length=new_length[:-1]
        try:
            #if the filtering goes wrong and less than 2 positive segments were found
            longest,second_longest=np.sort(new_length[new_value==1])[::-1][:2]
        except:
            print("less than 2 segments found at index{}".format(i))
            return np.nan,np.nan,np.nan,np.nan
        #finding the longest 2 segments which are far from centroid
        if np.where(np.logical_and(new_value==1,new_length==longest))[0].shape[0]==1:
            longest_positive_index=np.where(np.logical_and(new_value==1,new_length==longest))[0][0]
            second_longest_positive_index=np.where(np.logical_and(new_value==1,new_length==second_longest))[0][0]
        else:
            #in case the 2 segments are of equal length
            longest_positive_index=np.where(np.logical_and(new_value==1,new_length==longest))[0][0]
            second_longest_positive_index=np.where(np.logical_and(new_value==1,new_length==second_longest))[0][1]
        if longest_positive_index!=0:
            longest_positive_interval=[np.sum(length[:longest_positive_index]),np.sum(length[:longest_positive_index+1])-1]
        else:
            longest_positive_interval=[np.sum(length[:-1])%len(contour),length[0]-1]
        if second_longest_positive_index!=0:
            second_longest_positive_interval=[np.sum(length[:second_longest_positive_index]),np.sum(length[:second_longest_positive_index+1])-1]
        else:
            second_longest_positive_interval=[np.sum(length[:-1])%len(contour),length[0]-1]
        if (head_index<=longest_positive_interval[1] and 
            head_index>=longest_positive_interval[0]) or (head_index>=longest_positive_interval[0] 
            and longest_positive_interval[0]>longest_positive_interval[1]):
            #if head in the first segment,return the second segment
            return second_longest_positive_interval[0],second_longest_positive_interval[1],longest_positive_interval[0],longest_positive_interval[1]
        elif (head_index<=second_longest_positive_interval[1] and 
              #if head in the second segment,return the first segment
            head_index>=second_longest_positive_interval[0]) or (head_index>=second_longest_positive_interval[0] 
            and second_longest_positive_interval[0]>second_longest_positive_interval[1]):
            return longest_positive_interval[0],longest_positive_interval[1],second_longest_positive_interval[0],second_longest_positive_interval[1]
        else:
            #if head index not inside the valid segment when the fish is too curved?
            #choose tail segment as the segment furthur from the head_index
            dist_to_longest=min((head_index-longest_positive_interval[0])%len(contour),(head_index-longest_positive_interval[1])%len(contour),
                                (longest_positive_interval[0]-head_index)%len(contour),(longest_positive_interval[1]-head_index)%len(contour))
            dist_to_second_longest=min((head_index-second_longest_positive_interval[0])%len(contour),(head_index-second_longest_positive_interval[1])%len(contour),
                                (second_longest_positive_interval[0]-head_index)%len(contour),(second_longest_positive_interval[1]-head_index)%len(contour))
            if dist_to_second_longest>=dist_to_longest:
                return second_longest_positive_interval[0],second_longest_positive_interval[1],longest_positive_interval[0],longest_positive_interval[1]
            else:
                return longest_positive_interval[0],longest_positive_interval[1],second_longest_positive_interval[0],second_longest_positive_interval[1]
            
    tail_start,tail_end,head_start,head_end=find_segment(value,length)
    curviness_score_tail=compute_cos_fullbody(contour,step=step[0])
    curviness_score_head=compute_cos_fullbody(contour,step=step[1])
    curviness=curviness_score_tail[:,2]
    blurred_curviness=averageBlur(curviness,neighbor_width[0])

    if np.isnan(tail_start):
        return np.nan,np.nan,curviness_score_tail,length
    tail_segment=LoopingArray(blurred_curviness)[tail_start:tail_end+1]
    tail_index=(np.argmax(tail_segment)+tail_start)%l
    curviness=curviness_score_head[:,2]
    blurred_curviness=averageBlur(curviness,neighbor_width[1])
    head_segment=LoopingArray(blurred_curviness)[head_start:head_end+1]
    better_head_index=(np.argmax(head_segment)+head_start)%l
    return better_head_index,tail_index,curviness_score_tail,length

def compute_TailAngle_Dev(head_index,tail_index,contour):
    N=len(contour)
    head_midline=[contour[head_index%N],(contour[(head_index+50)%N]+contour[(head_index-50)%N])/2]
    tail_midline=[contour[tail_index%N],(contour[(tail_index+50)%N]+contour[(tail_index-50)%N])/2]
    vec1=np.array(head_midline[0]-head_midline[1])
    vec2=np.array(tail_midline[0]-tail_midline[1])
    cos=np.sum(vec1*vec2)/(np.linalg.norm(vec1)*np.linalg.norm(vec2))
    #numeric validity check
    cos=max(min(cos,1),-1)
    angle=np.arccos(cos)/np.pi*180
    headx1,heady1=head_midline[0]
    headx2,heady2=head_midline[1]
    tailx,taily=tail_midline[0]
    if headx1==headx2:
        d=abs(tailx-headx1)
        k1=10000
        b1=-k1*headx1
    elif heady1==heady2:
        d=abs(taily-heady1)
        k1=0
        b1=heady1
    else:
        k1=(heady1-heady2)/(headx1-headx2)
        b1=heady2-k1*headx2
        d=abs(k1*tailx-taily+b1)/np.sqrt(k1**2+1)
    return angle,d


class Args:
  video_path = None
  h5_path = None
  output_root = sys.argv[2]
  thres = 50 #thresholding parameter 0-255, 0 = black, 255 = white
  shrink_kernel = 3 #the parameter in cv2.erode and cv2.dilate


IMAGELENGTH=500
fps=40

args = Args()
args.video_path = os.path.join('/burg/zi/users/cpe2108/videos/', sys.argv[1])
args.h5_path = os.path.join('/burg/zi/users/cpe2108/h5/', str(sys.argv[1].split('.mp4')[0]) + 'DLC_resnet50_DLC_toptrackFeb27shuffle1_1030000.h5')
print(args.h5_path)
videopath = args.video_path
video_path = videopath
h5_path = args.h5_path
start = int(sys.argv[3])
stop = int(sys.argv[4])
fish_name = ".".join(os.path.basename(video_path).split(".")[:-1])
output_dir = os.path.join(args.output_root,fish_name)
if not os.path.exists(output_dir):
    os.makedirs(output_dir)

path = h5_path
f = pd.HDFStore(path,'r')
df = f.get('df_with_missing')
df.columns = df.columns.droplevel()
df=df.iloc[start:stop,:]
new_features=features(starttime=0,endtime=stop-start)
filtered_df=new_features.filter_df(df,add_midpoint=True)
filtered_head=df.A_head
filtered_head=filtered_head.fillna(method="ffill")
filtered_head=filtered_head.fillna(method="bfill") #just to make sure no NAs
new_features.fit(filtered_df)
features_df,_,_,_ = new_features.export_df()
features_df.to_csv(os.path.join(output_dir,"base_features.csv"),index = False)

#get a better contour using head position from DLC and thresholding again.
#the previous mask is also needed to remove some noise near fish's contour caused by raising the threshold
new_contour_array=find_contour(video_path,filtered_head,start=start,length = stop - start,step=1,threshold = args.thres,shrink_kernel = args.shrink_kernel,pre_filter = pre_filter)
with open(os.path.join(output_dir,"refined_contour"), "wb") as fp:
    pickle.dump(new_contour_array, fp)

decorated_result = cv2.VideoWriter(os.path.join(output_dir,"valid_video0.mp4"),cv2.VideoWriter_fourcc(*'mp4v'), fps, (IMAGELENGTH,IMAGELENGTH))
i = 0
cap = cv2.VideoCapture(videopath) 
index = start
cap.set(1,index)
while True:
    ret, frame = cap.read() 
    if not ret or index >= min(start+10000,stop):
        break
    if (index - start) %1000 == 0:
        print("current index {}".format(index))
        frame = cv2.cvtColor(frame,cv2.COLOR_BGR2RGB)

        cur_contour = new_contour_array[i]
        frame = cv2.drawContours(frame,[cur_contour],0,(255,0,0),2)
        decorated_result.write(frame)
        index += 1
        i += 1
decorated_result.release()
print("complete extract contour")  
#find head and tail position
curve_scores=[]
tail_indexs=[]
better_head_indexs=[]
lengths=[]
head_x=np.array(filtered_head.x)
head_y=np.array(filtered_head.y)
#no interpolation this time, contour coordinates should be within 0-500
for i in tqdm(range(len(new_contour_array))):
    contour=new_contour_array[i]
    head_index=head_on_contour(head_x[i], head_y[i], contour)
    better_head_index,tail_index,curve_score,length=predict_tail(contour,head_index,step=[33,57],neighbor_width=[17,17],minimal_seg_length=15)
    tail_indexs.append(tail_index)
    curve_scores.append(curve_score)
    lengths.append(length)
    better_head_indexs.append(better_head_index)

head_indexs=pd.Series(better_head_indexs).fillna(method="ffill").fillna(method = "bfill").astype(int)
tail_indexs=pd.Series(tail_indexs).fillna(method="ffill").fillna(method = "bfill").astype(int)
    
with open(os.path.join(output_dir,"curve_scores"), "wb") as fp:
    pickle.dump(curve_scores, fp)
    
with open(os.path.join(output_dir,"tail_index"), "wb") as fp:
    pickle.dump(tail_indexs.tolist(), fp)   
    
with open(os.path.join(output_dir,"head_index"), "wb") as fp:
    pickle.dump(head_indexs.tolist(), fp)    

with open(os.path.join(output_dir,"fish_segment_length"), "wb") as fp:
    pickle.dump(lengths, fp)       
    
print("tail feature extracted!")
tail_angles=[]
tail_devs=[]
for i in range(len(new_contour_array)):
    contour=curve_scores[i][:,:2]
    contour=contour.squeeze()
    head_index=head_indexs[i]
    tail_index=tail_indexs[i]
    tail_angle,tail_dev=compute_TailAngle_Dev(head_index,tail_index,contour)
    tail_angles.append(tail_angle)
    tail_devs.append(tail_dev)
tail_angles=np.array(tail_angles)
tail_devs=np.array(tail_devs)

with open(os.path.join(output_dir,"tailAngle"),"wb") as fp:
    pickle.dump(tail_angles, fp)    

with open(os.path.join(output_dir,"tail_dev"), "wb") as fp:
    pickle.dump(tail_devs, fp)    

#make validation video
decorated_result = cv2.VideoWriter(os.path.join(output_dir,"valid_video.mp4"),cv2.VideoWriter_fourcc(*'mp4v'), fps, (IMAGELENGTH,IMAGELENGTH))
i = 0
cap = cv2.VideoCapture(videopath) 
index = start
cap.set(1,index)
while True:
    ret, frame = cap.read() 
    if not ret or index >= min(start+10000,stop):
        break
    if (index - start) %1000 == 0:
        print("current index {}".format(index))
        frame = cv2.cvtColor(frame,cv2.COLOR_BGR2RGB)
        cur_contour = new_contour_array[i]
        cur_contour = np.array(cur_contour,dtype = np.int32)
        cur_tail_index = tail_indexs.iloc[i]
        l = len(cur_contour)
        head_x,head_y = filtered_head.x.iloc[i],filtered_head.y.iloc[i]
        tail_x,tail_y = new_contour_array[i][cur_tail_index%l][0]
        
        frame = cv2.circle(frame, (int(head_x),int(head_y)), 5, (0,0,255), -1)
        frame = cv2.circle(frame, (int(tail_x),int(tail_y)), 5, (0,255,0), -1)
        frame = cv2.drawContours(frame,[cur_contour],0,(255,0,0),2)
        decorated_result.write(frame)
        index += 1
        i += 1
decorated_result.release()
```
