## Supplemental File 2 for "Coordination and persistence of aggressive visual communication in Siamese fighting fish": contour_utils.py

```
#!/usr/bin/env python3
# -*- coding: utf-8 -*-
"""
Created on Mon Aug  3 14:29:42 2020

"""

import h5py
import numpy as np
import pandas as pd
import matplotlib.pyplot as plt
import seaborn as sn
import cv2
from moviepy.video.fx.all import crop
from moviepy.editor import VideoFileClip, VideoClip, clips_array
from moviepy.video.io.bindings import mplfig_to_npimage
from tqdm import tqdm
from scipy.ndimage import zoom  
import warnings
from scipy.sparse import coo_matrix
from LoopingArray import LoopingArray

IMAGELENGTH=500
fps=40

def relative_position_check(head,max_dist=20,max_counter=7):
    #try to keep the predicted position of head around the actual head as much as possible
    #assumptions: the first frame of head is accurate
                 #inaccurate head position(mostly be at tail) can last no more than 8 frames
                #if this happens, another 8 frames of mislocatedh head position will spawn
    #the reason it is not integrated in the class filtering function is because it is not done vector-wise and is likely to me much slower
    
    data = head.copy()
    head_x=data.x.iloc[0]
    head_y=data.y.iloc[0]
    assert ~np.isnan(head_x),"starting point should not be nan"
    counter=0
    for i in tqdm(range(data.shape[0])):
        x=data.x.iloc[i]
        y=data.y.iloc[i]
        distance=np.sqrt(np.square((x-head_x))+np.square((y-head_y)))
            #sometimes DLC misclassifies tail as head, so just to remove this by looking at the relative location changed
        if (distance>max_dist and counter<=max_counter) or np.isnan(distance) or np.isnan(x):
                #if cur point is close to the previous invalid pt, or cur point is far from prev valid pt
            data.x.iloc[i]=np.nan
            data.y.iloc[i]=np.nan
            counter+=1
        else:
            #record the latest valid_head position
            head_x=data.x.iloc[i]
            head_y=data.y.iloc[i]
            counter=0
    return data


#%%
#calculate the curvature at each point of the contour,default step=3

#currently only testing cosine angle
def compute_pointness(contour, n=3):
    out=np.zeros((600,600))
    contour=contour.squeeze()
    N = len(contour)
    t=1/N
    for i in range(N):
        x_cur, y_cur = contour[i]
        x_next, y_next = contour[(i + n) % N]
        x_prev, y_prev = contour[(i - n) % N]
        dy1=(y_next-y_prev)/(2*n*t)
        dx1=(x_next-x_prev)/(2*n*t)
        dy2=(y_next+y_prev-2*y_cur)/(n*n*t*t)
        dx2=(x_next+x_prev-2*x_cur)/(n*n*t*t)
        curvature=(dx1*dy2-dy1*dx2)/(np.power(dx1*dx1+dy1*dy1,3/2))
        curvature=abs(curvature)
        out[x_cur,y_cur]=curvature
    return out

#find the largest contour
def find_largest_contour(contours,interpolate=0):
    flag=0
    area=0
    for cnt in contours:
        new_area=cv2.contourArea(cnt)
        if new_area>area and new_area<10000*(interpolate*8+1) :
            area=new_area
            fish_contour=cnt
            flag=1
    if flag==1:
        return fish_contour,flag
    else:
        return np.zeros((1,1,2)),flag


def find_centroid(contour):
    moments=cv2.moments(contour)
    m01=moments.get("m01")
    m10=moments.get("m10")
    m00=moments.get("m00")
    xbar=m10/m00
    ybar=m01/m00
    return xbar,ybar

def compute_dist(contour,xbar,ybar):
    contour=contour.squeeze()
    dist=np.linalg.norm(contour-np.array([xbar,ybar]),axis=1)
    return dist


#give a heatmap along fish's contour about how curved it is
def plot_result(curvatures,contour,img_size=600,quantile=0.5,to_array=False):
    '''
    curvatures:the curvescore on the contour, should be an nxn array equalto img_size
    contour:one specific contour, nx1x2 array
    '''
    head_tail_x,head_tail_y=np.where(curvatures<0)
    new_curvatures=curvatures.copy()
    new_curvatures[new_curvatures<0]=0
    x,y=np.nonzero(new_curvatures)
    xbar,ybar=find_centroid(contour)
    colors=np.array(np.zeros_like(x),np.float64)
    dists=compute_dist(contour,xbar,ybar)
    for i in range(len(x)):
        colors[i]=new_curvatures[x[i],y[i]]
    fig =plt.figure()
    ax = fig.add_subplot(1,1,1)
    circ=plt.Circle((xbar, ybar), radius=np.quantile(dists,quantile),linewidth=0.5, color='red',fill=False)
    ax.add_patch(circ)
    plt.scatter(head_tail_x,head_tail_y,s=3,c="lemonchiffon")
    plt.scatter(x, y, c=colors, s=3,cmap="YlGnBu", vmin=0, vmax=0.2)#
    plt.plot(xbar,ybar,"ro",markersize=3,color="red")
    plt.xlim(0,img_size)
    plt.ylim(0,img_size)
    plt.colorbar()
    plt.title("curveness heatmap on fish contour")
    if not to_array:
        plt.show()
    else:
        out=mplfig_to_npimage(fig)
        plt.close(fig)
        return out

#exclude curvatures in head and tail
def filter_curvature(curvatures,contour):
    xbar,ybar=find_centroid(contour)
    dists=compute_dist(contour,xbar,ybar)
    quantile=np.quantile(dists,0.6)
    contour=contour.squeeze()
    N=len(contour)
    
    for i in range(N):
        x,y=contour[i]
        if dists[i]>quantile:
            curvatures[x,y]=0.1
    return curvatures


#find the 4 boundary pts of the contour
#instead of using local flipping, use run length encoding to determine the longest 4 segments
def find_anchor_rle(contour,validity=None,quantile=0.5,min_length=100,warning_cnt=0):
    '''
    parameters:
        contour:fish's contour, likely an nx1x2 array
        validity: a nx2 boolean array which tells whether the contour point is valid(not too close to head/tail),
        if not provided, it will be autogenerated from contour
        quantile: the threshold to remove head/tail parts, every pts having distance to centroid larger than quantile(dist)
        will be deemed invalid
        warning_cnt:for debugging, since the function will only output 4indexs, it shall cnt for the times where it actually
        will find 3 more segments
    '''
    contour=contour.squeeze()
    #if the valid pts are given before hand, skip this step
    if validity is None or quantile!=0.5:
        xbar,ybar=find_centroid(contour)
        dists=compute_dist(contour,xbar,ybar)
        thres=np.quantile(dists,quantile)
        validity=dists<thres
    value,length=runLengthEncoding(validity)
   #squueze the rle lists by combining those shorter segments
    def shrink_rle(value,length):
        #this also will just append those short segments to positive areas
        new_val=[]
        new_length=[]
        l=len(value)
        if value[0]!=value[l-1]:
            value.append(value[0])
            length.append(0)
        #if startpoint is actually at boundary, add a 0 length rle to it
        for i in range(l-1):
            if i==0:
                if (length[0]+length[l-1])>min_length:
                    new_val.append(value[0])
                else:
                    new_val.append(-1)
                new_length.append(length[0])
            elif length[i]<min_length:
                #prev element in new_val is unknown or 1
                if new_val[::-1][0]!=0:
                    ll=len(new_length)-1
                    new_length[ll]+=length[i]
                else:
                    #if value negative, save this length to the next segment
                    new_val.append(-1)
                    new_length.append(length[i])
            else:
                ll=len(new_length)-1
                if new_val[::-1][0]==-1:
                    if value[i]==0: #saved length for previous, because it starts from 0
                        #so the previous segment is actually 1
                        new_val[ll]=True
                        new_val.append(False)
                        new_length.append(length[i])
                    else:
                        #the cur long segment is 1, so concat the previous short segments
                        new_val[ll]=1
                        new_length[ll]+=length[i]
                else:
                    if new_val[::-1][0]!=value[i]:
                        new_val.append(value[i])
                        new_length.append(length[i])
                    else:
                        new_length[ll]+=length[i]
            #put the last segment in
        ll=len(new_length)-1
        if new_val[ll]==-1:
            new_val[ll]=1
        if (length[0]+length[l-1])>min_length:
            new_val.append(value[l-1])
            new_length.append(length[l-1])
        else:
            #if the whole segment in the start point is too short, just give it to the last valid segment
            new_length[ll]+=length[l-1]
        return new_val,new_length
    new_val,new_length=shrink_rle(value,length)
    if len(new_val)<=3:
        warning_cnt+=1
        print("\r"+"less than 4 segments detected, already happens {} times".format(warning_cnt),end="")
        return np.nan,np.nan,np.nan,np.nan,warning_cnt
    elif new_val[0]:
        out2=new_length[0]
        in1=new_length[1]+new_length[0]
        out1=new_length[2]+new_length[1]+new_length[0]
        in2=min(len(validity)-1,new_length[3]+new_length[2]+new_length[1]+new_length[0])
    else:
        in1=new_length[0]
        out1=new_length[1]+new_length[0]
        in2=new_length[2]+new_length[1]+new_length[0]
        out2=min(len(validity)-1,new_length[3]+new_length[2]+new_length[1]+new_length[0])
    return in1,out1,in2,out2,warning_cnt

#compute the cos angle of a point on contour to its previous pt and next pt
def compute_cos(contour, step=3,img_size=600,min_step=2,quantile=0.5):
    #I plan to return 3 objects :an img_size x img_size array with each point having the cosine angle value on it,
    #  2 lists which is the cosine in left and right part
    out=np.zeros((img_size,img_size))
    left_cosines=[]
    right_cosines=[]
    contour=contour.squeeze()
    xbar,ybar=find_centroid(contour)
    dists=compute_dist(contour,xbar,ybar)
    quantile=np.quantile(dists,quantile)
    validity=dists<quantile
    N = len(contour)
    t=1/N 
    def find_next(i,step,min_step):
        #find next point avaliable
        #say if index i+step is in the valid pts, then use i+step as tbe next point
        #otherwise backtrack the points until the point is valid or meet the minimal length requirement
        if step<=min_step or validity[(i+step)%N]==True:
            return contour[(i+step) % N]
        else:
            return find_next(i,step-1,min_step)
    def find_prev(i,step,min_step):
        if step<=min_step or validity[(i-step)%N]==True:
            return contour[(i -step) % N]
        else:
            return find_prev(i,step-1,min_step)
    for i in range(N):
        x_cur, y_cur = contour[i]
        if validity[i]==False:
            #head/tail position's cosine angle encoded to -1 for future visualization
            out[x_cur,y_cur]=-1
        else:
            x_next, y_next = find_next(i,step,min_step)
            x_prev, y_prev = find_prev(i,step,min_step)
            vec1=np.array([x_next-x_cur,y_next-y_cur])
            vec2=np.array([x_prev-x_cur,y_prev-y_cur])
            cos=np.sum(vec1*vec2)/np.sqrt(np.sum(vec1*vec1)*np.sum(vec2*vec2))
            out[x_cur,y_cur]=cos+1
    return out

  
#run length encoding:
#array: 1,0,0,0,1,1 -> length:1,3,2;value
def runLengthEncoding(arr): 
    value=[]
    length=[]
    prev=arr[0]
    value.append(prev)
    l=0
    for val in arr:
        if val==prev:
            l+=1
        else:
            length.append(l)
            value.append(val)
            l=1
        prev=val
    length.append(l)
    return np.array(value),np.array(length)


#find a conservative mask to exclude reflections and noises
def find_contour(videopath,head,length=40*300,start=0,step=1,pre_filter=None,threshold = 50,shrink_kernel = 3):
    '''
    prefilter:a function that changes mask array
    length: Total number of frames needed
    start: which frame to start
    step:pick every * frame
    thres: thresholding parameter when doing cv2.threshold
    shrink_kernel: how much you want to shrink the thresholded mask to avoid reflections
    '''
    vidcap = cv2.VideoCapture(videopath)
    contour_array=[]
    length=int(length/step)
    index=start
    vidcap.set(1,index)
    first = True
    for i in tqdm(range(length)):
        success,image=vidcap.read()
        if success!=1:
            print("process stops early at {}th iteration".format(i))
            break
        if pre_filter is not None:
            image = pre_filter(image)
        index+=step
        vidcap.set(1,index)
        gray = cv2.cvtColor(image, cv2.COLOR_BGR2GRAY)
        ret,th = cv2.threshold(gray, threshold, 255, cv2.THRESH_BINARY)
        if shrink_kernel>0:
          k2 = cv2.getStructuringElement(cv2.MORPH_RECT, (shrink_kernel,shrink_kernel))
          th = cv2.dilate(th, k2, iterations=1)
        #filter specific large black area, like 888 sign in other videos
        contours, hierarchy = cv2.findContours(np.invert(th.copy()), cv2.RETR_TREE, cv2.CHAIN_APPROX_SIMPLE)#simplified contour pts
        #finding the contour with largest area
        fish_contour,flag=find_largest_contour(contours)
        if flag==0:
            print("no valid fish contour find at index {}".format(i))
            img=np.float32(np.full(image.shape,0))#just in case there's no valid contour, won't happen in the current case
        else:
            #draw only the mask of this contour
            img=cv2.drawContours(np.float32(np.full(image.shape,255)),[fish_contour],0,(0,0,0),cv2.FILLED)
        img=cv2.cvtColor(img,cv2.COLOR_BGR2GRAY)
        if shrink_kernel>0:
          k3=cv2.getStructuringElement(cv2.MORPH_RECT, (shrink_kernel+2,shrink_kernel+2))
        else:
          k3=cv2.getStructuringElement(cv2.MORPH_RECT, (3,3))
        img=np.invert(np.array(cv2.erode(img,k3,iterations=1),np.uint8))
        #smoothing
        img=cv2.medianBlur(img, 5)
        true_contour, hierarchy = cv2.findContours(img.copy(), cv2.RETR_TREE, cv2.CHAIN_APPROX_NONE)#complete contour pts
        true_contour,flag=find_largest_contour(true_contour)
        if flag == 0:
          contour_array.append(np.random.randn(100,1,2))
          continue
        conservative_mask = np.array(cv2.drawContours(np.float32(np.full((img.shape[0],img.shape[1]),255)),[true_contour],0,(0,0,0),cv2.FILLED),np.uint8)
        conservative_contour = true_contour
        #re-thresholding image and try to include tail into the contour
        ret,th = cv2.threshold(gray, threshold, 255, cv2.THRESH_BINARY)
        k1 = cv2.getStructuringElement(cv2.MORPH_RECT, (5,5))
        th = cv2.erode(th, k1, iterations=1)

        xbar,ybar=find_centroid(conservative_contour)
        dists=compute_dist(conservative_contour,xbar,ybar)
        thres=np.quantile(dists,0.4)
        y,x=np.nonzero(np.invert(th))#black area, i.e. fish body
        head_x=head.x.iloc[i]
        head_y=head.y.iloc[i]
        th=exclusion(y,x,thres,xbar,ybar,head_x,head_y)           
        th=np.minimum(th,conservative_mask)
        th=np.invert(th)
        contours, hierarchy = cv2.findContours(th, cv2.RETR_TREE, cv2.CHAIN_APPROX_SIMPLE)
        fish_contour,flag=find_largest_contour(contours)
        if flag==0:
            print("no valid fish contour find at index {}".format(index-1))
            img=np.float32(np.full(image.shape,0))#just in case there's no valid contour, won't happen in the current case
        else:
            img=cv2.drawContours(np.float32(np.full(image.shape,255)),[fish_contour],0,(0,0,0),cv2.FILLED)
        img=cv2.cvtColor(img,cv2.COLOR_BGR2GRAY)
        #to get rid of small holes in the mask
        k3=cv2.getStructuringElement(cv2.MORPH_RECT, (3,3))
        img=np.invert(np.array(cv2.erode(img,k3,iterations=1),np.uint8))
        img=cv2.medianBlur(img, 5)
        true_contour, hierarchy = cv2.findContours(img.copy(), cv2.RETR_TREE, cv2.CHAIN_APPROX_NONE)#complete contour pts
        true_contour,_=find_largest_contour(true_contour,interpolate=1)
        try:
            contour_array.append(true_contour)
        except:
            contour_array.append(np.zeros((1,1,2),dtype=np.float64))
        
        if first: #plot sample contour 
            sample_image = cv2.drawContours(image.copy(),[true_contour],0,(255,0,0),2)
            plt.imshow(sample_image)
            plt.savefig("sample_contour.png")
            print("sample contour of the first frame has been saved to sample_contour.png")
            first = False
        
    return contour_array
def exclusion(y,x,thres,xbar,ybar,head_x,head_y):
    dists1=compute_dist(np.vstack([x,y]).T,xbar,ybar)
    #new found part should not be too close to the centroid(as it is tail)
    flag1=dists1<thres
    dists2=compute_dist(np.vstack([x,y]).T,head_x,head_y)
    #new found part shall not be too close to the head, otherwise it is likely a reflection
    flag2=dists2<thres
    flag=np.logical_or(flag1,flag2)
    x=x[~flag]
    y=y[~flag]
    data=np.full((len(x),),255)
    output=np.invert(np.uint8(coo_matrix((data,(y,x)),shape=(IMAGELENGTH,IMAGELENGTH)).toarray()))
    return output 

def head_on_contour(head_x,head_y,contour):
    dists=compute_dist(contour,head_x,head_y)
    index=np.argmin(dists)
    return index

def compute_cos_fullbody(contour, step=30):
    #compute the curvature for full body
    curviness=np.zeros((0,3),dtype=np.float64)
    contour=contour.squeeze()
    N = len(contour)
    def find_next(i,step):
        return contour[(i+step) % N]
    def find_prev(i,step):
        return contour[(i -step) % N]
    for i in range(N):
        x_cur, y_cur = contour[i]
        x_next, y_next = find_next(i,step)
        x_prev, y_prev = find_prev(i,step)
        vec1=np.array([x_next-x_cur,y_next-y_cur])
        vec2=np.array([x_prev-x_cur,y_prev-y_cur])
        cos=np.sum(vec1*vec2)/np.sqrt(np.sum(vec1*vec1)*np.sum(vec2*vec2))
        curviness=np.concatenate((curviness,np.array([x_cur,y_cur,cos+1]).reshape(1,3)))
    return curviness

def averageBlur(arr,neighbor_width=30):
    #updating an array of curviness, make every element to the avg value of its neighbor
    #So the tail tip is more likely to get a higher score compared to its neighbors
    out=arr.copy()
    l=len(arr)
    for i in range(l):
        out[i]=np.sum(LoopingArray(arr)[(i-neighbor_width):i+neighbor_width+1])/(2*neighbor_width+1)
    return out

#after the runlengthEncoding on validity derived from distance to centroid,
#some segments should be combined into longer segments
def combine_small_segment(value,length,minimal_length=70):
    l=len(value)
    flag=0
    #check if the starting point is a flipping point
    if value[0]==value[l-1]:
        temp=length[l-1]
        flag=1
        length[0]+=temp
        value=value[:l-1]
        length=length[:l-1]
    new_val=[]
    new_length=[]
    stack=0
    for i in range(len(value)):
        ll=length[i]
        #if the segment is smaller than certain length, put it to a existing segment
        #if there is no such thing, prepare that for a future valid segment
        if ll<minimal_length:
            if new_val:                
                new_length[len(new_length)-1]+=ll
            else:
                stack+=ll
        else:
            if len(new_val)==0 or new_val[::-1][0]!=value[i]:
                new_val.append(value[i])
                new_length.append(length[i]+stack)
                stack=0
            else:
                new_length[len(new_length)-1]+=ll
    if flag==1:
        #put the length back when we first cut off the last piece at first
        if new_val[len(new_length)-1]!=new_val[0]:
            new_val.append(new_val[0])
            new_length.append(temp)
        else:
            new_length[len(new_length)-1]+=temp
        new_length[0]-=temp
            
    return np.array(new_val),np.array(new_length)

def predict_tail(contour,head_index,step=None,quantile=0.4,neighbor_width=None,minimal_seg_length=23):
    #default value, in case the image is/is not zoomed
    if step is None:
        step=[int(len(contour)/10),int(len(contour)/10)*1.5]
    if neighbor_width is None:
        neighbor_width=[int(len(contour)/20),int(len(contour)/20)]
    xbar,ybar=find_centroid(contour)
    dists=compute_dist(contour,xbar,ybar)
    thres=np.quantile(dists,quantile)
    #get whether the pts are far from the centroid
    validity=dists>thres
    l=len(dists)
    value,length=runLengthEncoding(validity)
    value,length=combine_small_segment(value, length,minimal_seg_length)
    def find_segment(value,length):
        #index of segment
        l=len(value)
        new_length=length.copy()
        new_value=value.copy()
        #append the last segment unfinished segment to the front to compare the length
        if value[l-1]==value[0]:
            new_length[0]=length[0]+length[l-1]
            new_value=value[:-1]
            new_length=new_length[:-1]
        try:
            #if the filtering goes wrong and less than 2 positive segments were found
            longest,second_longest=np.sort(new_length[new_value==1])[::-1][:2]
        except:
            print("less than 2 segments found at index{}".format(i))
            return np.nan,np.nan,np.nan,np.nan
        #finding the longest 2 segments which are far from centroid
        if np.where(np.logical_and(new_value==1,new_length==longest))[0].shape[0]==1:
            longest_positive_index=np.where(np.logical_and(new_value==1,new_length==longest))[0][0]
            second_longest_positive_index=np.where(np.logical_and(new_value==1,new_length==second_longest))[0][0]
        else:
            #in case the 2 segments are of equal length
            longest_positive_index=np.where(np.logical_and(new_value==1,new_length==longest))[0][0]
            second_longest_positive_index=np.where(np.logical_and(new_value==1,new_length==second_longest))[0][1]
        if longest_positive_index!=0:
            longest_positive_interval=[np.sum(length[:longest_positive_index]),np.sum(length[:longest_positive_index+1])-1]
        else:
            longest_positive_interval=[np.sum(length[:-1])%len(contour),length[0]-1]
        if second_longest_positive_index!=0:
            second_longest_positive_interval=[np.sum(length[:second_longest_positive_index]),np.sum(length[:second_longest_positive_index+1])-1]
        else:
            second_longest_positive_interval=[np.sum(length[:-1])%len(contour),length[0]-1]
        if (head_index<=longest_positive_interval[1] and 
            head_index>=longest_positive_interval[0]) or (head_index>=longest_positive_interval[0] 
            and longest_positive_interval[0]>longest_positive_interval[1]):
            #if head in the first segment,return the second segment
            return second_longest_positive_interval[0],second_longest_positive_interval[1],longest_positive_interval[0],longest_positive_interval[1]
        elif (head_index<=second_longest_positive_interval[1] and 
              #if head in the second segment,return the first segment
            head_index>=second_longest_positive_interval[0]) or (head_index>=second_longest_positive_interval[0] 
            and second_longest_positive_interval[0]>second_longest_positive_interval[1]):
            return longest_positive_interval[0],longest_positive_interval[1],second_longest_positive_interval[0],second_longest_positive_interval[1]
        else:
            #if head index not inside the valid segment when the fish is too curved
            #choose tail segment as the segment furthur from the head_index
            dist_to_longest=min((head_index-longest_positive_interval[0])%len(contour),(head_index-longest_positive_interval[1])%len(contour),
                                (longest_positive_interval[0]-head_index)%len(contour),(longest_positive_interval[1]-head_index)%len(contour))
            dist_to_second_longest=min((head_index-second_longest_positive_interval[0])%len(contour),(head_index-second_longest_positive_interval[1])%len(contour),
                                (second_longest_positive_interval[0]-head_index)%len(contour),(second_longest_positive_interval[1]-head_index)%len(contour))
            if dist_to_second_longest>=dist_to_longest:
                return second_longest_positive_interval[0],second_longest_positive_interval[1],longest_positive_interval[0],longest_positive_interval[1]
            else:
                return longest_positive_interval[0],longest_positive_interval[1],second_longest_positive_interval[0],second_longest_positive_interval[1]
            
    tail_start,tail_end,head_start,head_end=find_segment(value,length)
    curviness_score_tail=compute_cos_fullbody(contour,step=step[0])
    curviness_score_head=compute_cos_fullbody(contour,step=step[1])
    curviness=curviness_score_tail[:,2]
    blurred_curviness=averageBlur(curviness,neighbor_width[0])

    if np.isnan(tail_start):
        return np.nan,np.nan,curviness_score_tail,length
    tail_segment=LoopingArray(blurred_curviness)[tail_start:tail_end+1]
    tail_index=(np.argmax(tail_segment)+tail_start)%l
    curviness=curviness_score_head[:,2]
    blurred_curviness=averageBlur(curviness,neighbor_width[1])
    head_segment=LoopingArray(blurred_curviness)[head_start:head_end+1]
    better_head_index=(np.argmax(head_segment)+head_start)%l
    return better_head_index,tail_index,curviness_score_tail,length

def compute_TailAngle_Dev(head_index,tail_index,contour):
    N=len(contour)
    head_midline=[contour[head_index%N],(contour[(head_index+50)%N]+contour[(head_index-50)%N])/2]
    tail_midline=[contour[tail_index%N],(contour[(tail_index+50)%N]+contour[(tail_index-50)%N])/2]
    vec1=np.array(head_midline[0]-head_midline[1])
    vec2=np.array(tail_midline[0]-tail_midline[1])
    cos=np.sum(vec1*vec2)/(np.linalg.norm(vec1)*np.linalg.norm(vec2))
    #numeric validity check
    cos=max(min(cos,1),-1)
    angle=np.arccos(cos)/np.pi*180
    headx1,heady1=head_midline[0]
    headx2,heady2=head_midline[1]
    #headx2,heady2=head_midline[1]
    tailx,taily=tail_midline[0]
    if headx1==headx2:
        d=abs(tailx-headx1)
        k1=10000
        b1=-k1*headx1
    elif heady1==heady2:
        d=abs(taily-heady1)
        k1=0
        b1=heady1
    else:
        k1=(heady1-heady2)/(headx1-headx2)
        b1=heady2-k1*headx2
        d=abs(k1*tailx-taily+b1)/np.sqrt(k1**2+1)
    return angle,d
```
