## Supplemental File 2 for "Coordination and persistence of aggressive visual communication in Siamese fighting fish": find_features.py

```
#!/usr/bin/env python3
# -*- coding: utf-8 -*-
"""
Created on Thu Jun  4 13:36:18 2020

"""

from functions import *
import pandas as pd
import numpy as np
import matplotlib.pyplot as plt
from scipy import interpolate 
from auto_filter_full import auto_scoring_tracefilter_full, transform_data,filter_tailbeating

class Feature_extraction():
    
    '''
    #extract feature from given period
    #function to compare the cluster outputs of different features, the visualization is ultilized on the scatter plot of orientation and operculum angle
    #requires filtered df has the same schema as that predefined filtered_df, which should contain column
    #'A_head',"F_spine1",'mid_spine1_spine2',"G_spine2",'mid_spine2_spine3',"H_spine3","I_spine4","J_spine5","K_spine6","L_spine7","B_rightoperculum",
    #'E_leftoperculum'
    '''
    
    def __init__(self,starttime=100000,endtime=None,duration=60):
        '''
        starttime: starttime of the period use
        duration: the duration of the period, the data will be sliced from starttime:starttime+40*duration
        '''
        self.starttime=starttime
        self.duration=duration
        if endtime==None:
            endtime=starttime+40*duration
        self.endtime=endtime
        
    def filter_df(self,raw_df,add_midpoint=True,p_head = 30,p_tail=15,angle_tol=75,p=0.5,p0=50,p1=25, p2 = 10, t1 = 20):
        '''
       
        Parameters
        ----------
        raw_df : TYPE 
            DESCRIPTION. Input raw DataFrame
        add_midpoint : TYPE, optional
            DESCRIPTION.  True:Yuyang's method, which will add 2 more columns(midpoint of spine1-spine2, and midpoint of spine2-spine3)
            False:Other method'

        Returns 
        -------
        filtered data
        '''

        if add_midpoint:
            df=transform_data(raw_df)
            return auto_scoring_tracefilter_full(df,p_head=p_head,p_tail=p_tail,p=p,angle_tol=angle_tol)
        else:
            return filter_tailbeating(raw_df,p0=p0,p_head = p_head, p1=p1, p2 = p2, t1 = t1)
            
        
    def fit(self,filtered_df,filter_feature=True,fill_na=True,estimate_na=True):
        '''
       this function computes all the features 
        Parameters
        ----------
        filtered_df : dataframe
            filtered dataframe
        filter_feature: filter out extreme points in the feature calculated, basically it removes points which violates the 
        3 /sigma rule
        fill_na: whether to fill na after the features are calculated
        estimate_na: whether to estimate the nas before fitting the spline, assuming the missing point is on the line of it's
        previous and next available point

        Returns 
        -------
      
        '''
        starttime=self.starttime
        endtime=self.endtime
        trunc_df=filtered_df.loc[starttime:endtime-1,:]
        operculum=auto_scoring_get_opdeg(trunc_df)
        
        ori=orientation(trunc_df)
    
        turn_angle=turning_angle_spine(trunc_df)
        
        mov_speed=speed(trunc_df)
        
        if "mid_spine1_spine2" in filtered_df.columns:
            y=trunc_df.loc[:,[('A_head','y'),("F_spine1","y"),('mid_spine1_spine2',"y"),("G_spine2","y"),
                            ('mid_spine2_spine3',"y"),("H_spine3","y"),("I_spine4","y"),("J_spine5","y"),
                            ("K_spine6","y"),("L_spine7","y")]]
            x=trunc_df.loc[:,[('A_head','x'),("F_spine1","x"),('mid_spine1_spine2',"x"),("G_spine2","x"),
                            ('mid_spine2_spine3',"x"),("H_spine3","x"),("I_spine4","x"),("J_spine5","x"),
                            ("K_spine6","x"),("L_spine7","x")]]
        else:
            y=trunc_df.loc[:,[('A_head','y'),("F_spine1","y"),("G_spine2","y"),
                            ("H_spine3","y"),("I_spine4","y"),("J_spine5","y"),
                            ("K_spine6","y"),("L_spine7","y")]]
            x=trunc_df.loc[:,[('A_head','x'),("F_spine1","x"),("G_spine2","x"),
                            ("H_spine3","x"),("I_spine4","x"),("J_spine5","x"),
                            ("K_spine6","x"),("L_spine7","x")]]
      
        concat_array=np.stack((x,y),axis=-1)
        if estimate_na:
            #give estimate to na's 
            concat_array=np.array(list(map(self.linearly_fill_data,concat_array)))
        map_results=np.array(list(map(self.cal_curvature_and_dir,concat_array)))
        curvatures=pd.DataFrame(map_results[:,0,:])
        cos=pd.DataFrame(map_results[:,1,:])
        diff_curvature=pd.DataFrame(np.diff(curvatures,axis=0,prepend=np.expand_dims(curvatures.loc[0,:],0)))
        if "mid_spine1_spine2" in filtered_df.columns:
            curvatures.columns=["curvature_head","curvature_spine1","curvature_spine1.5","curvature_spine2","curvature_spine2.5","curvature_spine3",
              "curvature_spine4","curvature_spine5","curvature_spine6","curvature_spine7"]
            diff_curvature.columns=["diff_curvature_head","diff_curvature_spine1","diff_curvature_spine1.5","diff_curvature_spine2","diff_curvature_spine2.5","diff_curvature_spine3",
              "diff_curvature_spine4","diff_curvature_spine5","diff_curvature_spine6","diff_curvature_spine7"]
            cos.columns=["tangent_head","tangent_spine1","tangent_spine1.5","tangent_spine2","tangent_spine2.5","tangent_spine3",
              "tangent_spine4","tangent_spine5","tangent_spine6","tangent_spine7"]
        else:
            curvatures.columns=["curvature_head","curvature_spine1","curvature_spine2","curvature_spine3",
              "curvature_spine4","curvature_spine5","curvature_spine6","curvature_spine7"]
            diff_curvature.columns=["diff_curvature_head","diff_curvature_spine1","diff_curvature_spine2","diff_curvature_spine3",
              "diff_curvature_spine4","diff_curvature_spine5","diff_curvature_spine6","diff_curvature_spine7"]
            cos.columns=["tangent_head","tangent_spine1","tangent_spine2","tangent_spine3",
              "tangent_spine4","tangent_spine5","tangent_spine6","tangent_spine7"]
        if filter_feature:
            #filter feature according to "3 sigma" rule, that is, I assume the features follows normal distribution, and 
            # I filter out points where it's distance to its mean is greater than 3xstd
            operculum[abs(operculum-np.nanmean(operculum,axis=0))>3*np.nanstd(operculum,axis=0)]=np.nan
            ori[abs(ori-np.nanmean(ori,axis=0))>3*np.nanstd(ori,axis=0)]=np.nan
            turn_angle[abs(turn_angle-np.nanmean(turn_angle,axis=0))>3*np.nanstd(turn_angle,axis=0)]=np.nan
            mov_speed[abs(mov_speed-np.nanmean(mov_speed,axis=0))>3*np.nanstd(mov_speed,axis=0)]=np.nan
            curvatures[abs(curvatures-np.nanmean(curvatures,axis=0))>3*np.nanstd(curvatures,axis=0)]=np.nan
            diff_curvature[abs(diff_curvature-np.nanmean(diff_curvature,axis=0))>3*np.nanstd(diff_curvature,axis=0)]=np.nan
            cos[abs(cos-np.nanmean(cos,axis=0))>3*np.nanstd(cos,axis=0)]=np.nan
            
        #then deal with the NAs in the feature
        if fill_na==True:       
            curvatures=curvatures.fillna(method='ffill'); curvatures=curvatures.fillna(curvatures.mean())
            cos=cos.fillna(method='ffill').fillna(cos.mean())
            diff_curvature=diff_curvature.fillna(method='ffill'); diff_curvature=diff_curvature.fillna(diff_curvature.mean())
            operculum=operculum.fillna(method="ffill").fillna(operculum.mean())
            ori=pd.Series(ori).fillna(method='ffill').fillna(np.nanmean(ori))
            mov_speed=pd.Series(mov_speed).fillna(method="ffill").fillna(np.nanmean(mov_speed))
            turn_angle=pd.Series(turn_angle).fillna(method="ffill").fillna(np.nanmean(turn_angle))
        self.curvatures=curvatures
        self.cos=cos
        self.curvatures=curvatures
        self.diff_curvature=diff_curvature
        self.operculum=operculum
        self.ori=ori
        self.mov_speed=mov_speed
        self.turn_angle=turn_angle
    
    def linearly_fill_data(self,x):
        ##Yuqi's code, filter step is skipped, I will just show na in the curvatures computed
        not_na = np.unique(np.where(~np.isnan(x))[0])
        if (len(not_na)>=4):
            h = not_na[0]
            s1 = not_na[1]
            if (h == 0) & (s1==1):
                for j in range(len(not_na)):
                    if j > 1:
                        current = not_na[j]
                        pre = not_na[j-1]
                        point = current-pre
                        if point > 1:
                            #if there's point missing in consective samples for spline,fill the missing points in 
                            #the middle with a linear estimate
                            dx = x[current][0]-x[pre][0]
                            dy = x[current][1]-x[pre][1]
                            for k in range(1, point):
                                x[pre+k][0] = x[pre][0]+k*dx/point
                                x[pre+k][1] = x[pre][1]+k*dy/point
        return x

    def cal_curvature_and_dir(self,x):
        pts=x
        line=pts[2]-pts[1]
        index=~np.isnan(pts).any(axis=1)
        pts=pts[index]
        curvature=np.repeat(np.nan,x.shape[0])
        directions=np.repeat(np.nan,x.shape[0])
        if(pts.shape[0]>=4):
            tck,u=interpolate.splprep(pts.T, u=None, s=0.0)
            dx1,dy1=interpolate.splev(u,tck,der=1)
            dx2,dy2=interpolate.splev(u,tck,der=2)
            k=(dx1*dy2-dy1*dx2)/np.power((np.square(dx1)+np.square(dy1)),3/2)
            direction=(dy1*line[1]+dx1*line[0])/np.linalg.norm(line)/np.sqrt(dy1*dy1+dx1*dx1)
            directions[index]=direction
            curvature[index]=k
        return [curvature,directions]
    
    def export_df(self):
        #combine curvature/diff_curvature/tangent_angle and other features to one df
        other_features=pd.DataFrame({"operculum":np.array(self.operculum),"orientation":self.ori,"movement_speed":np.array(self.mov_speed),
                                     "turning_angle":self.turn_angle},index=self.curvatures.index)
        curvatures=self.curvatures
        diff_curvatures=self.diff_curvature
        tangent=self.cos
        return other_features,curvatures,diff_curvatures,tangent
        
    def visualize_cluster(self,num_cluster=2,dpi=300,s=2,cmap='cividis'):
        '''
        dpi: the resolution of image?
        num_cluster: the number of cluster in kmeans
        s:size of pts
        cmap:cmap attribute in plt
        '''
        #scale them
        from sklearn.preprocessing import StandardScaler
        operculum=self.operculum;ori=self.ori
        filled_curvatures=self.curvatures;diff_curvature=self.diff_curvature;filled_cos=self.cos
        scaler = StandardScaler();scaler.fit(filled_curvatures);filled_curvatures=pd.DataFrame(scaler.transform(filled_curvatures))   
        scaler = StandardScaler();scaler.fit(diff_curvature); diff_curvature=pd.DataFrame(scaler.transform(diff_curvature))
        scaler = StandardScaler();scaler.fit(filled_cos);filled_cos=pd.DataFrame(scaler.transform(filled_cos)) 
        #kmeans cluster, probably not the optimal way to do this
        from sklearn.cluster import KMeans
        kmeans_curvature = KMeans(n_clusters=num_cluster, init='k-means++', max_iter=1000, n_init=10);kmeans_curvature.fit(filled_curvatures)
        kmeans_diffCurvature = KMeans(n_clusters=num_cluster, init='k-means++', max_iter=1000, n_init=10); kmeans_diffCurvature.fit(diff_curvature)
        kmeans_cos = KMeans(n_clusters=num_cluster, init='k-means++', max_iter=1000, n_init=10);kmeans_cos.fit(filled_cos)
    
        #visualize
        #it's not the most solid way to do this, just want to check the features don't look uniform distributed in the plot
        label_curvature=kmeans_curvature.predict(filled_curvatures)
        label_diffCurvature=kmeans_diffCurvature.predict(diff_curvature)
        label_cos=kmeans_cos.predict(filled_cos)
        self.label_curvature=label_curvature
        self.label_diffCurvature=label_diffCurvature
        self.label_cos=label_cos
        fig=plt.figure(dpi=dpi)
        ax=fig.add_subplot(1,3,1)
        ax.title.set_text("curvature")
        ax.scatter(x=operculum,y=ori,s=s, c=label_curvature, cmap='cividis')
        ax=fig.add_subplot(1,3,2)
        ax.scatter(x=operculum,y=ori,s=s, c=label_diffCurvature, cmap='cividis')
        ax.title.set_text("diff_curvature")
        ax=fig.add_subplot(1,3,3)
        ax.scatter(x=operculum,y=ori,s=s, c=label_cos, cmap='cividis')
        ax.title.set_text("tangent line")
        print("cluster on {} groups".format(num_cluster))

##The old name
class features(Feature_extraction):
    def  __init__(self,starttime=100000,endtime=None,duration=6):
        super(features,self).__init__(starttime,endtime,duration)
```
