## Supplemental File 2 for "Coordination and persistence of aggressive visual communication in Siamese fighting fish": functions.py

```
#!/usr/bin/env python3
# -*- coding: utf-8 -*-
"""
Created on Wed Mar  4 22:15:45 2020

"""

import numpy as np
import pandas as pd
from glob import glob
import os
from scipy.optimize import linear_sum_assignment
from scipy import optimize

def nanarccos (floatfraction):
    con1 = ~np.isnan(floatfraction)
    
    if con1:
        cos = np.arccos(floatfraction)
        OPdeg = np.degrees(cos)
    
    else:
        OPdeg = np.nan
    
    return (OPdeg)

def vecnanarccos():
    
    A = np.frompyfunc(nanarccos, 1, 1)
    
    return A
def lawofcosines(line_1,line_2,line_3):
    '''
    Takes 3 series, and finds the angle made by line 1 and 2 using the law of cosine
    '''
 
    num = line_1**2 + line_2**2 - line_3**2
    denom = (line_1*line_2)*2
    floatnum = num.astype(float)
    floatdenom = denom.astype(float)
    floatfraction = floatnum/floatdenom
    OPdeg = vecnanarccos()(floatfraction)
    
    return OPdeg

def mydistance(pos_1,pos_2):
    '''
    Takes two position tuples in the form of (x,y) coordinates and returns the distance between two points
    '''
    x0,y0 = pos_1
    x1,y1 = pos_2
    dist = np.sqrt((x1-x0)**2 + (y1-y0)**2)

    return dist

def coords(data):
    '''
    helper function for more readability/bookkeeping
    '''
    return (data['x'],data['y'])

def auto_scoring_get_opdeg(data_auto):
    '''
    Function to automatically score operculum as open or closed based on threshold parameters. 
    
    Parameters: 
    data_auto: traces of behavior collected as a pandas array. 
    thresh_param0: lower threshold for operculum angle
    thresh_param1: upper threshold for operculum angle
    
    Returns:
    pandas array: binary array of open/closed scoring
    '''
    # First collect all parts of interest:
    poi = ['A_head','B_rightoperculum','E_leftoperculum']
    HROP = mydistance(coords(data_auto[poi[0]]),coords(data_auto[poi[1]]))
    HLOP = mydistance(coords(data_auto[poi[0]]),coords(data_auto[poi[2]]))
    RLOP = mydistance(coords(data_auto[poi[1]]),coords(data_auto[poi[2]]))
    
    Operangle = lawofcosines(HROP,HLOP,RLOP)
    
    return Operangle

def getfiltereddata(h5_files):
    ''' takes dlc files and returns 2 filtered dataframes. Helpful when doing quick analysis of two opponents and 
    there is a need to flip one of the opponent's coordinates. Flipping coordinates is needed to align certain metrics
    like orientation whether a fish is on teh right side facing left stimulus or left side facing right stimulus. '''
    file_handle1 = h5_files[0]

    with pd.HDFStore(file_handle1,'r') as help1:
        data_auto1 = help1.get('df_with_missing')
        data_auto1.columns= data_auto1.columns.droplevel()
        data_auto1_filt = auto_scoring_tracefilter (data_auto1)
     
    file_handle2 = h5_files[1]

    with pd.HDFStore(file_handle2,'r') as help2:
        data_auto2 = help2.get('df_with_missing')
        data_auto2.columns= data_auto2.columns.droplevel()
        data_auto2_filt = auto_scoring_tracefilter(data_auto2)
        data_auto2_filt['A_head']['x'] = data_auto2_filt['A_head']['x'] + 500
        data_auto2_filt['B_rightoperculum']['x'] = data_auto2_filt['B_rightoperculum']['x'] + 500
        data_auto2_filt['E_leftoperculum']['x'] = data_auto2_filt['E_leftoperculum']['x'] + 500
    
    return data_auto1_filt,data_auto2_filt


def auto_scoring_tracefilter(data,p0=20,p2=15):
    ''' takes dataframe and filters keypoints on the body based on how far point jumps between frames and how close point is to origin (0,0)'''

    mydata = data.copy()
    boi = ['A_head','B_rightoperculum', 'C_tailbase', 'D_tailtip','E_leftoperculum']
    for b in boi:
        for j in ['x','y']:
            xdifference = abs(mydata[b][j].diff())
            xdiff_check = xdifference > p0     
            mydata[b][j][xdiff_check] = np.nan
 
            origin_check = mydata[b][j] < p2
            mydata[origin_check] = np.nan

    return mydata
def gaze_tracking(fish1,fish2):
    """
    A function that takes in two dataframes corresponding to the two fish. Assymetric. Fish one is the one gazing, fish two is being gazed at. 
    
    Parameters: 
    Fish1: A dataframe of tracked points. 
    Fish2: A dataframe of tracked points. 

    Output:
    A vector of angles between 0 and 180 degrees (180 corresponds to directed gaze).
    """


    ## Get the midpoint of the gazing fish. 
    midx,midy = midpoint(fish1['B_rightoperculum']['x'], fish1['B_rightoperculum']['y'], fish1['E_leftoperculum']['x'], fish1['E_leftoperculum']['y'] )
    line1 = mydistance((midx, midy), (fish1['A_head']['x'], fish1['A_head']['y']))
    line2 = mydistance((fish1['A_head']['x'], fish1['A_head']['y']), (fish2['A_head']['x'], fish2['A_head']['y']))
    line3 = mydistance((fish2['A_head']['x'], fish2['A_head']['y']), (midx, midy))

    angle = lawofcosines(line1, line2, line3)
    return angle

def midpoint (pos_1, pos_2, pos_3, pos_4):
    '''
    give definition pos_1: x-value object 1, pos_2: y-value object 1, pos_3: x-value object 2
    pos_4: y-value object 2
    '''
    midpointx = (pos_1 + pos_3)/2
    midpointy = (pos_2 + pos_4)/2

    return (midpointx, midpointy)

def orientation2(data_auto):
    ## take out the head coordinate
    head_x = data_auto["A_head"]["x"]
    head_y = data_auto["A_head"]["y"]
    
    ## get the midpoint coordinate
    mid_x = midpoint(data_auto['B_rightoperculum']['x'], data_auto['B_rightoperculum']['y'], data_auto['E_leftoperculum']['x'], data_auto['E_leftoperculum']['y'])[0]
    mid_y = midpoint(data_auto['B_rightoperculum']['x'], data_auto['B_rightoperculum']['y'], data_auto['E_leftoperculum']['x'], data_auto['E_leftoperculum']['y'])[1]
    
    ## calculate cos theta
    cos_angle = list()
    for i  in range(data_auto.shape[0]):
        inner_product = (head_x[i]-mid_x[i])*head_x[i]
        len_product = ((head_x[i]-mid_x[i])**2+((head_y[i]-mid_y[i])**2))**(0.5)*head_x[i]
        cos_angle.append(inner_product/len_product)
    
    cos_angle = np.array(cos_angle)
    
    return cos_angle

def speed (data, fps = 40,n_prev=10):
    
    ''' function that calculates velocity of x/y coordinates
    plug in the xcords, ycords, relevant dataframe, fps
    return the velocity as column in relevant dataframe. unit output = frames/sec'''

    poi = ['A_head']
    (Xcoords, Ycoords)= coords(data[poi[0]])
    n_after_x=np.concatenate((np.repeat(np.nan,n_prev),Xcoords[:-n_prev]))
    n_after_y=np.concatenate((np.repeat(np.nan,n_prev),Ycoords[:-n_prev]))
    distx=n_after_x-Xcoords
    disty=n_after_y-Ycoords
    TotalDist = np.sqrt(distx**2 + disty**2)
    Speed = TotalDist / (n_prev/fps) #converts to seconds
#    
    return Speed

def periculum_speed(periculum,fps=40):
    periculum=periculum.fillna(method="bfill")
    movement=np.diff(periculum)
    movement=np.insert(movement,0,np.nan)
    p_speed=abs(movement/(1/fps))
    return p_speed
''' fit a sine curve for the opericulum data'''

def sin_curve(x, b,c,d):
    return  30*np.sin(b * x+c)+d
def find_params(periculum,start=100000,end=100400):
    params, params_covariance = optimize.curve_fit(sin_curve, range(start,end), periculum[start:end],p0=[0.05,0,70])
    return params

def find_error(periculum):
    params=find_params(periculum)
    predict=sin_curve(range(len(periculum)),params[0],params[1],params[2])
    error=predict-periculum
    return abs(error)

def turning_angle(data_auto):
    head_x = data_auto["A_head"]["x"]
    head_y = data_auto["A_head"]["y"]
    
    ## get the midpoint coordinate
    mid_x = midpoint(data_auto['B_rightoperculum']['x'], data_auto['B_rightoperculum']['y'], data_auto['E_leftoperculum']['x'], data_auto['E_leftoperculum']['y'])[0]
    mid_y = midpoint(data_auto['B_rightoperculum']['x'], data_auto['B_rightoperculum']['y'], data_auto['E_leftoperculum']['x'], data_auto['E_leftoperculum']['y'])[1]

    cur_vec=np.vstack((head_x-mid_x,head_y-mid_y)).T
    prev_vec=np.vstack((np.repeat(np.nan,40),np.append((head_x-mid_x)[:-40]),np.append(np.repeat(np.nan,40),(head_y-mid_y)[40:]))).T
    inner_product=np.sum(np.multiply(cur_vec,prev_vec),axis=1)
    cur_norm=np.sum(np.multiply(cur_vec,cur_vec),axis=1)
    prev_norm=np.sum(np.multiply(prev_vec,prev_vec),axis=1)
    cos=inner_product/np.sqrt(cur_norm*prev_norm)
    angle=np.arccos(cos)
    return angle/np.pi*180
def turning_angle_spine(data_auto):
    spine1_x = data_auto["F_spine1"]["x"]
    spine1_y = data_auto["F_spine1"]["y"]
    
    if "mid_spine1_spine2" in data_auto.columns:
        spine1_5_x=data_auto["mid_spine1_spine2"]["x"]
        spine1_5_y=data_auto["mid_spine1_spine2"]["y"]
    else:
        spine1_5_x=data_auto["G_spine2"]["x"]
        spine1_5_y=data_auto["G_spine2"]["y"]
    ##find the direction in the previous second
    cur_vec=np.vstack((spine1_x-spine1_5_x,spine1_y-spine1_5_y)).T
    prev_vec=np.vstack((np.append(np.repeat(np.nan,40),(spine1_x-spine1_5_x)[:-40]),np.append(np.repeat(np.nan,40),(spine1_y-spine1_5_y)[:-40]))).T
    inner_product=np.sum(np.multiply(cur_vec,prev_vec),axis=1)
    cur_norm=np.sum(np.multiply(cur_vec,cur_vec),axis=1)
    prev_norm=np.sum(np.multiply(prev_vec,prev_vec),axis=1)
    cos=inner_product/np.sqrt(cur_norm*prev_norm)
    angle=np.arccos(cos)
    return angle/np.pi*180

def find_local_max_per(periculum,width=20):
    '''use for loop, tedious'''
    re=[]
    for i in range(len(periculum)):
        local=periculum[max(0,i-width):min(len(periculum)-1,i+width)]
        re.append(local.max())
    return re
    
def orientation(data_auto):
    ## take out the head coordinate
    head_x = data_auto["A_head"]["x"]
    head_y = data_auto["A_head"]["y"]
    
    ## get the midpoint coordinate
    mid_x = midpoint(data_auto['B_rightoperculum']['x'], data_auto['B_rightoperculum']['y'], data_auto['E_leftoperculum']['x'], data_auto['E_leftoperculum']['y'])[0]
    mid_y = midpoint(data_auto['B_rightoperculum']['x'], data_auto['B_rightoperculum']['y'], data_auto['E_leftoperculum']['x'], data_auto['E_leftoperculum']['y'])[1]
    
    ## calculate cos theta
    head_ori=np.array([head_x-mid_x,head_y-mid_y]).T
    ref=np.array([1,0])
    inner_product =head_ori.dot(ref)
    cos=inner_product/np.sqrt(np.sum(np.multiply(head_ori,head_ori),axis=1))
    angle=np.arccos(cos)/np.pi*180

    return angle
#%%
def compute_cos(x,y):
    #x,y is a (n,2) vector, output a nx1 vector of cosine for each row
    inner_product=np.sum(np.multiply(x,y),axis=1)
    norm_x=np.sum(np.multiply(x,x),axis=1)
    norm_y=np.sum(np.multiply(y,y),axis=1)
    inner_product/np.sqrt(norm_x*norm_y)

def manual_scoring(data_manual,period_length,crop0 = 0,crop1= -1):
    '''
    A function that takes manually scored data and converts it to a binary array. 
    
    Parameters: 
    data_manual: manual scored data
    data_auto: automatically scored data, just used here to establish how long the session is. 
    
    Returns: 
    pandas array: binary array of open/closed scoring
    '''
    Manual = pd.DataFrame(0, index=np.arange(period_length), columns = ['OpOpen'])
    reference = data_manual.index
    
    
    for i in reference:
        Manual[data_manual['Start'][i]:data_manual['Stop'][i]] = 1
    
    print(Manual[data_manual['Start'][i]:data_manual['Stop'][i]]) 
     
    return Manual['OpOpen'][crop0:crop1]

'''
This is similar to the find_permutation function in ssm package, the only change is that I changed
the elements of the cost_matrix to be the distance between clusters
'''

def compute_state_overlap(z1, z2, K1=None, K2=None):
    assert z1.dtype == int and z2.dtype == int
    assert z1.shape == z2.shape
    assert z1.min() >= 0 and z2.min() >= 0

    K1 = z1.max() + 1 if K1 is None else K1
    K2 = z2.max() + 1 if K2 is None else K2

    overlap = np.zeros((K1, K2))
    for k1 in range(K1):
        for k2 in range(K2):
            overlap[k1, k2] = np.sum((z1 == k1) & (z2 == k2))
    return overlap


def find_permutation(z1, z2, K1=None, K2=None):
    overlap = compute_state_overlap(z1, z2, K1=K1, K2=K2)
    K1, K2 = overlap.shape

    tmp, perm = linear_sum_assignment(-overlap)
    assert np.all(tmp == np.arange(K1)), 

    # Pad permutation if K1 < K2
    if K1 < K2:
        unused = np.array(list(set(np.arange(K2)) - set(perm)))
        perm = np.concatenate((perm, unused))

    return perm

def permute(x,perms):
    #permute_labels
    re=x.copy()
    K=np.max(x)+1
    for i in range(K):
        re[x==i]=perms[i]
    return re
```
