## Supplemental File 2 for "Coordination and persistence of aggressive visual communication in Siamese fighting fish": LoopingArray.py

```
#!/usr/bin/env python3
# -*- coding: utf-8 -*-
"""
Created on Thu Aug 13 14:18:36 2020

"""
import numpy as np
class LoopingArray():
    #1D looping array
    def __init__(self,array):
        self.data = np.array(array)

    def __getitem__(self, key):
        l=len(self.data)
        if isinstance(key,int):
            #key is an integer
            return self.data[key%l]
        elif isinstance(key,slice):
            #key is a slice
            start=key.start
            stop=key.stop
            step=key.step
            if step==0:
                raise ValueError("step cannot be 0")
            if step is None:
                step=1   
            if start is None:
                if step>0:
                    start=0
                else:
                    start=l-1
            if stop is None:
                if step>0:
                    stop=l
                else:
                    stop=-1
            if (stop-start)*step<0:
                #assuming the index length will not exceed 2 times the length of the loopingArray
                stop+=l*int(step/abs(step))
            shape=list(self.data.shape)
            shape[0]=len(range(start,stop,step))
            re=np.zeros(shape)
            for ind,i in enumerate(range(start,stop,step)):
                re[ind]=self.data[i%l]
            return re
        else:
            raise TypeError('Invalid argument type: {}'.format(type(key)))
    def __setitem__(self, key, value):
        l=len(self.data)
        if isinstance(key,int):
            self.data[key%l] = value
        elif isinstance(key,slice):
            start=key.start
            stop=key.stop
            step=key.step
            if step==0:
                raise ValueError("step cannot be 0")
            if step is None:
                step=1
            if start is None:
                if step>0:
                    start=0
                else:
                    start=l-1
            if stop is None:
                if step>0:
                    stop=l
                else:
                    stop=-1
            if (stop-start)*step<0:
                #assuming the index length will not exceed 2 times the length of the loopingArray
                stop+=l*int(step/abs(step))
            for ind,i in enumerate(range(start,stop,step)):
                self.data[i%l]=value[ind]
        else:
            raise TypeError('Invalid argument type: {}'.format(type(key)))

    def __repr__(self):
        if self.data.shape[0]==1:
            return "LoopingArray({})".format(self.data)
        else:
            return "LoopingArray(\n{})".format(self.data)

    def __len__(self):
        return len(self.data)


   
```
