## Supplemental File 2 for "Coordination and persistence of aggressive visual communication in Siamese fighting fish": README.md

The main script "contour_auto_single_FINALT.py" calls functions from all other scripts:
- auto_filter_full.py
- contour_utils.py
- find_features.py
- functions.py
- LoopingArray.py

The script requires that two main inputs share the same beginning of the basename: 
1) Video (.mp4) (video_path); on line 375 (ex. test3.mp4)
2) DLC output file (.h5) (h5_path); on line 376 (ex. test3DLC_resnet50_DLC_toptrackFeb27shuffle1_1030000.h5)

The script takes 4 arguments: 
1) path to video file
2) path to desired output location (output root)
3) start_index
4) stop_index 

The script can thus be run with a terminal command in this structure:  

python3 <path to contour_auto_single_FINALT.py> <path to video file> <path to desired output location (output root)> start_index stop_index

Note that the specific calibrations of the contour dilation/blur have all been optimized for our specific recording set up and animals. Customization
steps may need to be applied to accomodate recordings with animals for instance closer to the camera or in different lighting conditions. However, these functions can act as guides. 
